# A new cheese population in *Penicillium roqueforti* and adaptation of the five populations to their ecological niche

**DOI:** 10.1101/2023.01.21.524518

**Authors:** Ewen Crequer, Jeanne Ropars, Jean-Luc Jany, Thibault Caron, Monika Coton, Alodie Snirc, Jean-Philippe Vernadet, Antoine Branca, Tatiana Giraud, Emmanuel Coton

**Affiliations:** Univ. Brest, INRAE, Laboratoire Universitaire de Biodiversité et Ecologie Microbienne, F-29280 Plouzané, France; Université Paris-Saclay, CNRS, AgroParisTech, Laboratoire Ecologie Systématique et Evolution, UMR 8079, Bâtiment 680, 12 route RD128, 91190 Gif-sur-Yvette, France

**Author notes:** These authors co-supervised the study (Tatiana Giraud, Emmanuel Coton).

**Keywords:** domestication, fungus, physiology, growth modelling, carbon sources, fungicide tolerance

## Abstract

Domestication is an excellent case study for understanding adaptation and multiple fungal lineages have been domesticated for fermenting food products. Studying domestication in fungi has thus both fundamental and applied interest. Genomic studies have revealed the existence of four populations within the blue-cheese-making fungus *Penicillium roqueforti*. The two cheese populations show footprints of domestication, but the adaptation of the two non-cheese populations to their ecological niches (*i.e*. silage/spoiled food and lumber/spoiled food) has not been investigated yet. Here, we reveal the existence of a new *P. roqueforti* population, specific to French Termignon cheeses, produced using small-scale traditional practices, with spontaneous blue mould colonisation. This Termignon population is genetically differentiated from the four previously identified populations, providing a novel source of genetic diversity for cheese making. Phenotypically, the non-Roquefort cheese population was the most differentiated, with specific traits beneficial for cheese making, in particular higher tolerance to salt, to acidic pH and to lactic acid. Our results support the view that this clonal population, used for many cheese types in multiple countries, is a domesticated lineage on which humans exerted strong selection. The Termignon population displayed substantial genetic diversity, both mating types, horizontally transferred regions previously detected in the non-Roquefort population, and intermediate phenotypes between cheese and non-cheese populations. The lumber/spoiled food and silage/spoiled food populations were not more tolerant to crop fungicides but showed faster growth in various carbon sources (*e.g*. dextrose, pectin, sucrose, xylose and/or lactose), which can be beneficial in their ecological niches. Such contrasted phenotypes between *P. roqueforti* populations, with beneficial traits for cheese-making in the cheese populations and enhanced ability to metabolise sugars in the lumber/spoiled food population, support the inference of domestication in cheese fungi and more generally of adaptation to anthropized environments.

## Introduction

Domestication, for which humans operate strong selection for identified traits in a species, has been used since Charles Darwin’s “*On the origin of species*” book (1859) as a model case of adaptive divergence. Adaptive divergence under domestication has been documented in various lineages in animals and plants (*e.g*. evolution of dogs from wolves and corn from teosinte; Steensels et al. 2019), but also in bacteria and fungi (*e.g*. the lactic acid bacteria *Oenococcus oeni*, Lorentzen et al. 2019, and the yeast *Saccharomyces cerevisiae*, Pontes et al. 2020). Adaptive divergence can result in highly differentiated domesticated varieties. For example, the spread and adaptation of maize to different agro-ecological and cultural environments has led to a multitude of varieties (Meyer & Purugganan, 2013). In dogs, the extreme behavioural and morphological differences between the 500 breeds are the result of directional selection for different behaviours, aspects and usages (Hayward *et al*., 2019). The magnitude of morphological differences between dog breeds is higher than that between wild Canidae (Wayne, 1986).

Fungi represent an interesting clade to study adaptive divergence as they exhibit small genomes and short generation times compared to most plant and animal models. Ascomycota (one of the two main groups of fungi, encompassing most moulds and yeasts) are particularly used for fermented food, to produce a variety of beverages and food from animal products (*e.g*. cheese and dry cured sausages; Venturini Copetti 2019) or plant products (*e.g*. wine, bread, tempeh and soya sauce; Nout et al. 2004). Among domesticated Ascomycota, *S. cerevisiae*, the baker’s yeast, has been extensively studied (Gallone *et al*., 2016; García-Ríos *et al*., 2017; Legras *et al*., 2018; Peter *et al*., 2018). Adaptive divergence has been demonstrated in *S. cerevisiae*, with population subdivision and differences between populations in terms of phenotypes relevant for fermentation, such as sulfite resistance, maltotriose fermentation and growth at low temperature (Gallone et al. 2016; García-Ríos et al. 2017). Moulds in Ascomycota have also been domesticated for fermenting food products. For example, *Aspergillus oryzae* (Gibbons *et al*., 2012; Watarai *et al*., 2019) and *Aspergillus sojae* (Chang, 2004), intensively used in Asia as koji moulds for the production of various fermented products (*e.g*. miso, sake and shoyu), have evolved beneficial traits for fermentation and human consumption, such as starch breakdown ability (Gibbons *et al*., 2012) and the lack of toxin production (Kiyota et al. 2011). *Penicillium camemberti*, the fungus used for making bloomy rind soft cheeses (*e.g*. Brie and Camembert), has also been domesticated. This clonal lineage is indeed genetically differentiated from non-cheese closely related populations and exhibits beneficial traits for cheese production, for example in terms of colour, growth rate, competitive ability and reduced mycotoxin production (Gillot et al., 2017; Ropars et al. 2020).

Another filamentous ascomycete fungus of interest for the study of adaptive divergence is *Penicillium roqueforti*, because of its importance in several countries for making different types of renowned blue cheeses (*e.g*. Roquefort, Fourme d’Ambert, Gorgonzola, Danish blue, Stilton and Cabrales). Within *P. roqueforti*, four genetically differentiated populations have been identified so far (Dumas *et al*., 2020), one population being associated with mouldy silage and spoiled food, one with lumber and spoiled food, and two different cheese populations, resulting from independent domestication events. One of the cheese populations, called non-Roquefort, corresponds to a clonal lineage (Dumas *et al*., 2020) largely used for the production of many blue cheeses in various countries (*e.g*. Gorgonzola, Cabrales, Danish blue and Stilton), but not for Roquefort cheeses produced under the protected designation of origin (PDO) label. Strains belonging to the non-Roquefort population harbour large horizontally transferred genomic regions, in particular the *CheesyTer* and *Wallaby* genomic regions (Cheeseman et al. 2014; Ropars et al. 2015), conferring advantages in terms of growth and competitive abilities (Ropars et al. 2015; Dumas et al 2020). These two horizontally transferred genomic regions are absent from the other three identified *P. roqueforti* populations but have also been acquired, completely or partially, by other cheese-making *Penicillium* fungi *(e.g. Penicillium camemberti* and *P. biforme*; Cheeseman et al. 2014; Ropars et al 2015). The non-Roquefort lineage has acquired multiple beneficial traits for commercial cheese production, very different from non-cheese populations, such as faster growth on cheese, better exclusion of spoilers, the ability to make bluer cheeses and more attractive aromas (Gillot *et al*., 2017; Coton *et al*., 2020; Dumas *et al*., 2020; Caron *et al*., 2021).

The other known cheese population is strongly associated with the Roquefort protected designation of origin (PDO; Gillot et al. 2015; Dumas et al. 2020). This Roquefort population is differentiated from the non-Roquefort population and displays higher genetic and phenotypic diversities (Gillot *et al*., 2017; Dumas *et al*., 2020), which is probably due to the requirement to use local strains for the Roquefort PDO (Cahier des charges de l’appellation d’origine “Roquefort”, 2017). This has prevented Roquefort-cheese producers from using the clonal non-Roquefort lineage, so that they kept using their strains originating from different cultures on local farms in the area. The Roquefort population also displays beneficial traits for cheese production, including the production of typical aromas and blue veined cheeses with larger blue areas than the non-cheese populations (Dumas et al. 2020; Caron et al. 2021). The Roquefort population however seems more adapted to ancient production modes, with for example greater spore production on bread, the original medium used to produce *P. roqueforti* conidia for cheese inoculation (Dumas et al. 2020). The Roquefort population also displays slower growth, which was likely beneficial for cheese production before refrigeration, to preserve cheeses over a span of several months (Dumas et al. 2020).

*Penicillium roqueforti* cheese strains analysed so far all came from cheeses inoculated with spores from cultivated strains. However, some local blue cheeses can be spontaneously colonised by *P. roqueforti*. This is the case for “Bleu de Termignon” cheeses produced by only a handful of cheese makers in the French Alps, with smallscale production. Spontaneous colonisation could result from the presence of feral strains in the farm environment or ripening caves, *i.e*. spores released from cheeses, corresponding to a known or unknown domesticated cheese population. Alternatively, the strains spontaneously colonising Termignon cheeses could correspond to a genuinely wild population. However, Roquefort-like cheeses produced with the non-cheese populations described so far produced blue cheeses with little or unpleasant aromas (Caron *et al*., 2021), pointing more to a feral origin or a locally domesticated population.

In addition to the cheese populations, two other populations have indeed been identified in *P. roqueforti*, one largely associated with silage and spoiled food (*e.g*., bread, fruits and jam) and the other mainly found in lumber and spoiled food (*e.g*., inner fridge wall and mouldy wild apple; Dumas et al. 2020). These non-cheese populations display higher levels of sexual fertility than the cheese populations and grow faster under harsh conditions (Ropars *et al*., 2016, 2020; Dumas *et al*., 2020). The non-cheese populations also present higher genetic diversities than the cheese populations and have diverged from each other more recently than their common ancestor from the cheese populations (Dumas *et al*., 2020). They may therefore correspond either to populations recently adapted to human-made environments different from cheeses or to ancestral-like, wild populations.

The occurrence of genetically differentiated populations in distinct ecological niches suggests adaptive divergence in *P. roqueforti*. Indeed, these ecological niches are either plant- or dairy-related and vary in terms of environmental conditions (*e.g*., temperature, pH and salt concentration) and composition (nutrients). Such contrasting environments likely require different metabolic activities for a fungus to thrive in. For example, temperature is much lower during cheese ripening (8 to 14°C according to the (Cahier des charges de l’appellation d’origine « Bleu d’Auvergne », 2016; Cahier des charges de l’appellation d’origine « Bleu du Vercors-Sassenage », 2017)) than for optimal silage production (20-30°C; Weinberg et al. 2001; Borreani and Tabacco 2010; Ferrero et al. 2021). pH in Roquefort cheeses is around 4.7 and ranges from 3.7 to 5 in silage depending on the forage type (*e.g*., maize or grass) and on the dry matter percent of the crop. Blue cheeses are among the cheeses with highest salt concentrations (Hashem *et al*., 2014). Beyond contributing to the organoleptic qualities as a taste enhancer, salt prevents contamination by undesired microorganisms, as it impairs both fungal and bacterial growth by reducing water availability (also called water activity, *a_w_*). Water availability is under regulation for cheese import in the USA, for consumer safety, as it prevents the growth of pathogenic microorganisms such as *Listeria* spp. Salt represents nearly 4% weight in Roquefort, leading to very low water activity values (Caron *et al*., 2021). For non-cheese food, the specific temperature, pH and water activity depend on the type of food and storage conditions. Cheese and silage are environments with high organic acid concentrations due to fermentation, with in particular large quantities of lactic acid (Bevilacqua & Califano, 1989). Lactic acid may be used as a carbon source by some microorganisms (Eschrich *et al*., 2002; Jiang *et al*., 2014; Schink *et al*., 2022) but also exhibits antimicrobial activities, being therefore commonly used as a food preservative.

Carbon sources are very different between the ecological niches of the distinct *P. roqueforti* populations. For example, maltose, starch, pectine, cellobiose and xylose are associated with plant material, while the sugars in cheese are lactose and galactose, but they are present only at the beginning of cheese making and mostly metabolised by bacteria (Lee *et al*., 2014). Some of the *P. roqueforti* ecological niches can include fungal inhibitors or fungicides; for example, the two *P. roqueforti* populations occurring on spoiled food may be exposed to chemical preservatives commonly used for food preservation (*e.g*. potassium sorbate and natamycin). In *P. roqueforti*, sorbate resistance has been reported in some strains isolated from spoiled food, which has been attributed to the presence of a horizontally transferred region named *SORBUS* (Punt et al. 2022). In addition, the *P. roqueforti* population associated with silage may be exposed to triazoles, commonly used as fungicides on crops. The existence of specific adaptation to silage and food ecological niches by their *P. roqueforti* populations has not been investigated so far.

In order to improve our understanding of adaptive divergence in *P. roqueforti*, we investigated here i) the genetic differentiation of strains isolated from Termignon blue cheeses from other populations, by comparing their genomes to available genomes of strains from other ecological niches, and by looking for three horizontally transferred regions *(Wallaby, CheesyTer* and *SORBUS*) previously found in some *P. roqueforti* populations or strains, and ii) the phenotypic differentiation between *P. roqueforti* populations for traits likely under differential selection in the different ecological niches. We studied the impact of temperature, pH, salt concentration, various carbon sources and fungal inhibitors on the growth rate and latency of the different *P. roqueforti* populations using laser nephelometry, a mid-throughput method which measures fungal growth by estimating particle density in liquid media based on the detection of light reflection by cells. We thereby tested the hypothesis that the Termignon cheeses are made with a specific *P. roqueforti* population, that has evolved adaptive traits for cheese making, and, more generally, that the different *P. roqueforti*populations exhibit adaptation to their respective ecological niches.

## Material and methods

### Strain collection and conidia suspension preparation

For comparative genomics, we analysed the genomes of 51 *P. roqueforti* strains (Suppl. Table 1). Among them, we randomly selected for phenotypic experiments seven strains from each of the non-Roquefort and Roquefort populations, six strains from the lumber/spoiled food population, and eight strains from the silage/spoiled food population, which presents the highest genetic diversity (Dumas *et al*., 2020). We also used the four available strains sampled from Termignon blue cheeses. All strains are part of the ESE or UBOCC (https://nouveau.univ-brest.fr/ubocc/fr) culture collections (Suppl. Table 1).

Conidia suspensions were prepared for the different experiments by cultivating the fungal strains for six days at 25°C on potato dextrose agar (PDA, Difco, Fisher Scientific). Two mL of Tween 80 (0.045 %, v/v) were then added on each plate and conidia were scraped off the surface. Conidia concentrations in the suspensions were estimated using Malassez cells and adjusted to 10^6^ conidia.mL^-1^ with Tween 80. Fresh suspensions were prepared for each experiment.

### Genome sequencing and population genomics

DNA was extracted for Termignon cheese isolates and all new strains sequenced using the Illumina technology for this project from fresh haploid mycelium and conidia after monospore isolation and growth for five days on malt agar using the Nucleospin Soil Kit (Macherey-Nagel). Sequencing was performed using the Illumina paired-end technology (Illumina Inc.) at the INRAe GenoToul and CNRS I2BC platforms. We also generated an improved long read-based genome assembly of the LCP06133 strain. Its DNA was extracted from mycelium and conidia with the NucleoBond High Molecular Weight DNA kit (Macherey-Nagel), with the mechanical disruption of about 30 mg of lyophilized mycelium with beads for 5 min at 30 Hz. Its genome was sequenced using the Oxford Nanopore MinION technology with an R9 flow cell without multiplexing. The Nanopore library was prepared with the SQK-LSK109 ligation sequencing kit, and sequencing was performed in-house. We assessed the run quality using Porechop version 0.2.3_sequan2.1.1 (Wick et al., 2017). All raw data are available at NCBI under the project IDXX (*to be completed upon manuscript acceptance)*. We also used some available genomes (Dumas *et al*., 2020) (Suppl. Table 1).

We trimmed all Illumina reads and cleaned adapters with Trimmomatic v0.36 (Bolger *et al*., 2014). We removed the three bases at the beginning and end of reads using the *Leading* and *Trailing* options, dropped reads having a length inferior to 36 with the *MINLEN* option and performed a sliding window trimming approach using the *Slidingwindow* option and 4:25 as values. Cleaned reads were mapped on the new high-quality *P. roqueforti* LCP06133 reference genome assembly (ID Genbank, *to be completed upon manuscript acceptance)* using *Bowtie2* version 2.3.4.1 (Langmead & Salzberg, 2012). This reference genome was built using raw ONT reads with Canu version 1.8 (Koren *et al*., 2017) with the option genomeSize=28m and polished twice using Illumina reads (Dumas *et al*., 2020) with Pilon version 1.24 (Walker *et al*., 2014)), after a mapping using *Bowtie2* (Langmead & Salzberg, 2012) with a maximum length (-X) of 1000bp for both runs. Redundant contigs were removed based on a self-alignment. In Bowtie2, the maximum length (-X) was set to 1000 and the preset “very-sensitive-local” was used. SAMtools v1.7 (Li *et al*., 2009) was used to filter out duplicate reads and reads with a mapping quality score above ten for SNP calling. Single nucleotide polymorphisms (SNPs) were called using the GATK v4.1.2.0 Haplotype Caller (McKenna *et al*., 2010), generating a gVCF file per strain. GVCFs were combined using GATK CombineGVCFs, genotypes with GATK GenotypeGVCFs, and SNPs were selected using GATK SelectVariants. SNPs were filtered using GATK VariantFiltration and options QUAL <30, DP < 10, QD < 2.0, FS > 60.0, MQ < 40.0, SOR > 3.0, QRankSum < −12.5, ReadPosRankSum < −8.0. All processes from cleaning to variant calling were performed with Snakemake v5.3.0 (Koster & Rahmann, 2012) using an internally developed script.

We used Splitstree v4.16.2 (Huson & Bryant, 2006) for the neighbour-net analysis. We used the R package *Ade4* (Chessel *et al*., 2004; Dray & Dufour, 2007; Bougeard & Dray, 2018; Thioulouse *et al*., 2018) for principal component analyses (PCA, centred and unscaled). We used NGSadmix v.33 (Jørsboe *et al*., 2017) from the ANGSD (Korneliussen *et al*., 2014) package (version 0.933-110-g6921bc6) to infer individual ancestry from genotype likelihoods based on realigned reads, by assuming a given number of populations. A Beagle file was first prepared from bam using ANGSD with the following parameters: “-uniqueOnly 1 -remove_bads 1 - only_proper_pairs 1 - GL 1-doMajorMinor 1 - doMaf 1 - doGlf 2-SNP_pval 1e-6”. The Beagle file was used to run NGSadmix with 4 as the minimum number of informative individuals. The analysis was run for different *K*values, ranging from 2 to 6. A hundred independent runs were carried out for each number of clusters (*K*).

The nucleotide diversity *π* (Nei’s Pi; Nei and Li 1979; Hudson et al. 1992), the Watterson’s *θ* (Watterson, 1975), the fixation index *F_ST_* (Hudson et al. 1992) and the absolute divergence *d_XY_* (Nei & Li, 1979) were calculated using the *popgenome* package in R (Pfeifer *et al*., 2014). Fixed, private and shared sites were counted using custom scripts available at https://github.com/BastienBennetot/fixed_shared_private_count, with bcftools version 1.11 (using htslib 1.13+ds). The strain ESE00421 was not included for these computations. The pairwise homology index (PHI) test was performed using Splitstree v4.16.2 (Huson & Bryant, 2006).

Mating types in the Termignon population were determined using a mapping approach: the two mating-type alleles previously identified in *P. roqueforti* (Ropars *et al*., 2012) were used as references to map the Termignon strain reads.

We assessed the presence of the horizontally transferred regions *Wallaby* and *CheesyTer* in the genomes of the Termignon strains (as they were previously searched for in the other genomes), and of the *SORBUS* region in all the genomes analysed here, by mapping the Illumina reads on the published sequences (Cheeseman *et al*., 2014; Ropars *et al*., 2015; Punt *et al*., 2022).

### Growth media and conditions for phenotypic studies

For all tested conditions, we used potato dextrose broth (PDB, Difco, Fisher Scientific), except to evaluate carbon source impact on growth, for which we used a minimum medium (see below). For assessing temperature impact on growth, microplates were incubated at nine different temperatures (4, 8, 12, 20, 22, 25, 27, 30 and 32°C). For evaluating the pH impact on growth, PDB (initial pH 5.2) was adjusted to the targeted pH values (2, 3, 4, 5, 6, 7, 8, 9, 10, 12 or 14) using either hydrochloric acid or sodium hydroxide. To assess the effect of salt concentration on growth, we added NaCl (Sigma-Aldrich, Germany), using eleven concentrations (1, 1.5, 2, 2.5, 3, 3.5, 4, 5, 6, 8 and 10%), corresponding to the following of water activity values (fraction of water available for growth, *a_w_*): 0.996, 0.991, 0.988, 0.986, 0.983, 0.980, 0.977, 0.974, 0.967, 0.960, 0.946 and 0.929. To evaluate the impact of lactic acid on growth, this acid was added to PDB to obtain the target concentrations (0.05, 0.1, 0.2, 0.3, 0.4, 0.5, 0.6, 0.8, 1 and 1.5 M) and pH was adjusted to 5.2 ± 0.1 with sodium hydroxide. All media were sterilised at 121°C for 15 minutes and stored at 4°C until use, except for pH-modified media which were instead filtered at 0.45 μm to avoid pH change during autoclaving. To study the impact of carbon sources on growth, we supplemented the “M0” minimum medium (urea: 0.5 g.L^-1^, magnesium sulphate: 0.25 g.L^-1^, biotin and thiamin: 50 mg.L^-1^, citric acid and zinc sulphate: 5 mg.L^-1^, iron alum: 1 mg.L^-1^, copper sulphate: 0.25 mg.L^-1^ manganese sulphate, boric acid, sodium molybdate: 0.05 mg.L^-1^) with 20 g.L^-1^ of either sucrose, glucose, lactose, galactose, maltose, cellobiose, xylose, starch, pectin or sodium lactate. The media were then sterilised at 121°C for 15 minutes. To assess the impact of fungicides on growth, we supplemented PDB with either potassium sorbate (Sigma-Aldrich, Germany, 0.5 or 1 g.L^-1^), natamycin (Sigma-Aldrich, Germany, 2 or 5 mg.L^-1^) or tebuconazole (Sigma-Aldrich, Germany, 2.5, 5 or 10 mg.L^-1^). We sterilised PDB 2X at 121°C for 15 minutes and inhibitor solutions by filtration at 0.45 μm, and then combined PDB 2X, inhibitor filtrate and sterile water in order to obtain the targeted inhibitor concentrations.

### Growth monitoring by laser nephelometry

We added 190 μL of broth culture media in sterile 96-well microplates (Thermo Scientific) and completed each well with either 10 μL of conidia suspension at 10^6^ conidia.mL^-1^ or with 10 μL of Tween 80 (0.045% v/v) for negative controls. For incubation, microplates were stored in a plastic box containing sterile distilled water to regulate humidity during incubation. Measures were obtained using a laser nephelometer (NEPHELOstar^PLUS^, BMG labtech) with a laser diode at 635 nm. At the beginning of each measurement, a 20-second 500 rpm double-orbital agitation step was performed by the nephelometer microplate reader (Savary et al. 2022). Each microplate was monitored four times a day for days 1-2, thrice a day for days 3-7, twice a day for days 8-14 and once a day for days 15-21 or until the signal reached saturation. For each experiment, a minimum of three replicas per strain and condition were performed for more robust estimates of growth parameters.

### Fungal growth kinetic follow-up and modelling

Growth kinetics were obtained with nephelometric measurement, *i.e*. measurement of forward scattered light that is directly proportional to the suspended particle concentration (*i.e*. the turbidity) and to mycelium dry matter (Joubert et al. 2010), see Suppl. Fig. S1. The results are expressed in an arbitrary unit named relative nephelometry unit (RNU which can range from zero to several thousands until detector saturation), corresponding to the measure of the scattered light at an angle relative to the incident light source to avoid possible transmitted light interference, as a function of time. In order to estimate growth latency and growth rate for each strain and parameter tested, we performed a primary modelling step with the MatLab software (R2018b, Natick, Massachusetts: The MathWorks Inc.; see Figure 2A). We fitted a growth curve to the measured values for each strain and tested parameter using the modified Gompertz equation (1) (Zwietering *et al*., 1990), as it is the most robust for fungal growth modelling (Declerck *et al*., 2001; Savary *et al*., 2022).

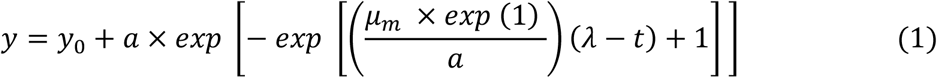

where *y_0_* is the value for time 0, *y* is the turbidity signal (in RNU) measured at time *t*, *a* is the maximal amplitude of the turbidity signal in RNU, *μ_m_* is the growth rate (RNU.h^-1^) in the exponential phase (hereafter called “maximal growth rate” as typically done in modelling growth studies) and *λ* is the lag time in hours *(i.e*. the time before fungal growth is detected). See Suppl. Fig. S1 for an illustration of the parameters.

**Figure 1:**
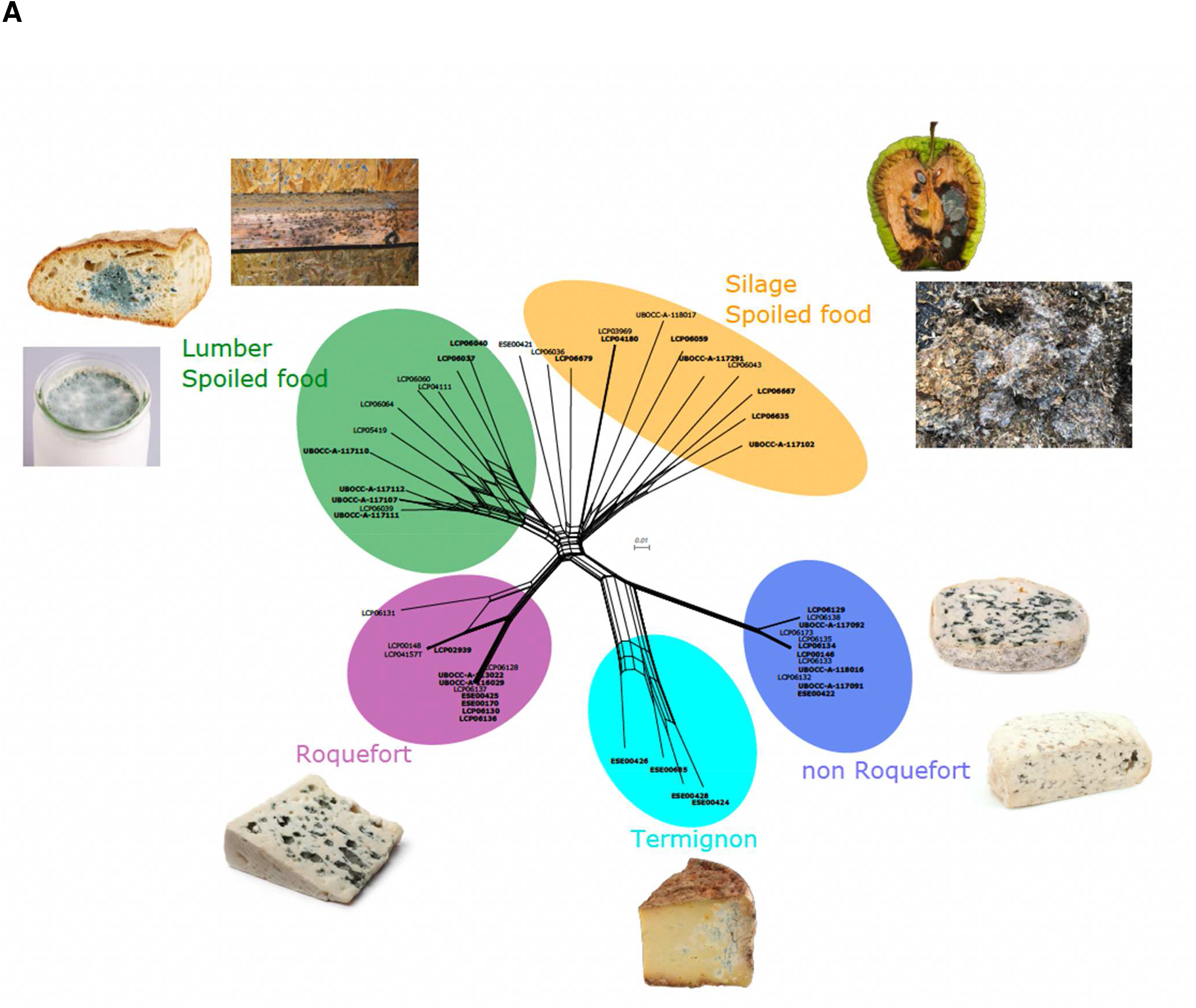
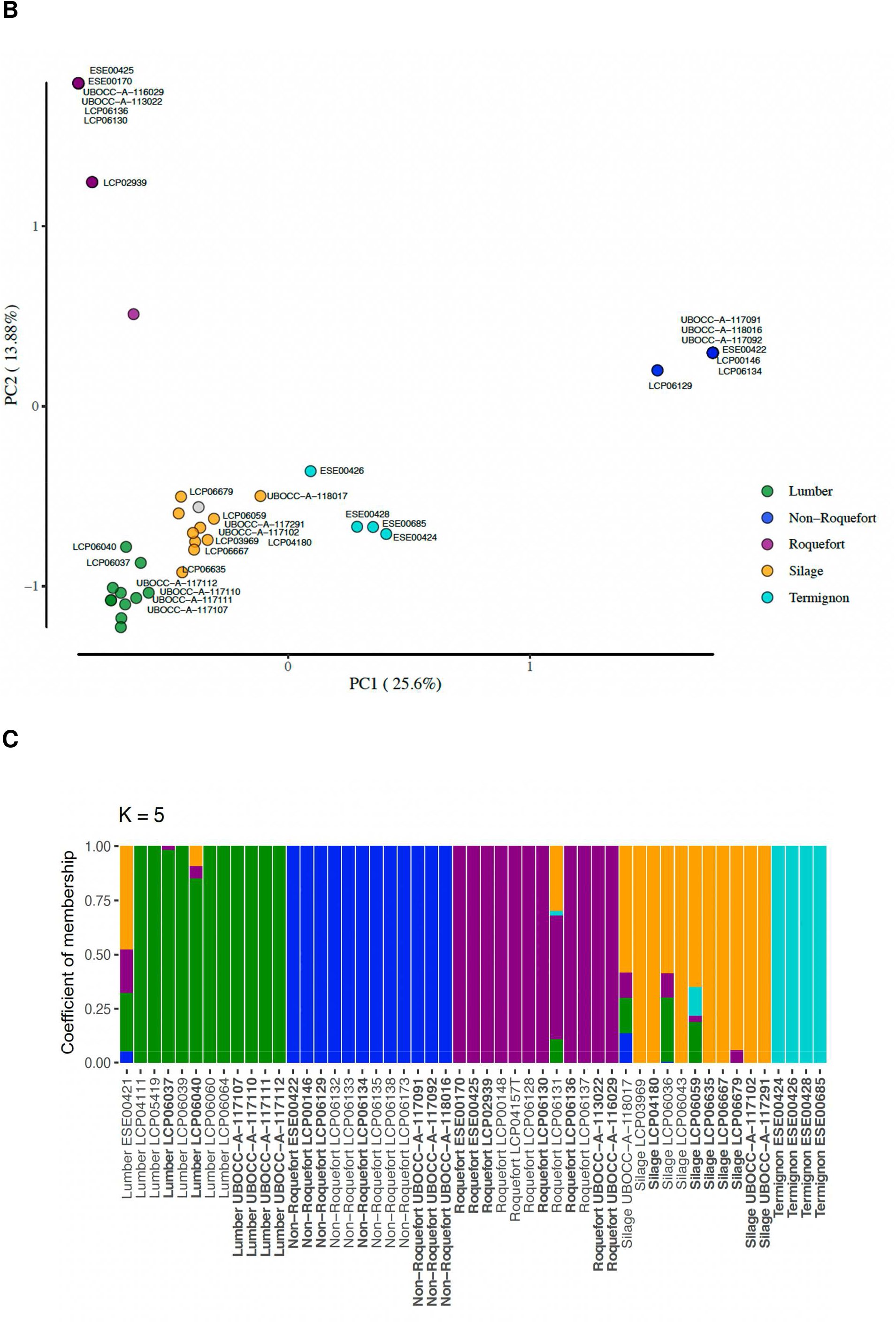
Representation of the genomic data and the differentiation between the five populations of *Penicillium roqueforti:* the three cheese populations (Roquefort, non-Roquefort and Termignon) and the two non-cheese populations (silage/food spoiler and lumber/spoiled food). **A.** Reticulated network of *P. roqueforti* strains based on 190,387 single nucleotide polymorphisms, showing the five distinct populations with pictures of their respective environments of collection. The ID of the strains used for phenotyping are in bold. **B.** Principal component analysis based on 190,387 single nucleotide polymorphisms. The names of the strains used in phenotype comparisons are indicated. The strain ESE00421 with intermediate assignments in various clusters with NGSadmix is shown in grey. **C**. Population subdivision inferred with NGSadmix for K=5 populations (See Suppl. Fig. 1 for K=2 to 6). Colored bars represent the coefficients of membership in the K gene pools based on genomic data. Each bar represents a strain, its name being indicated at the bottom of the figure. The ID of the strains used for phenotyping are in bold. The same colour code as on the other figures is used in all three panels.

**Figure 2:**
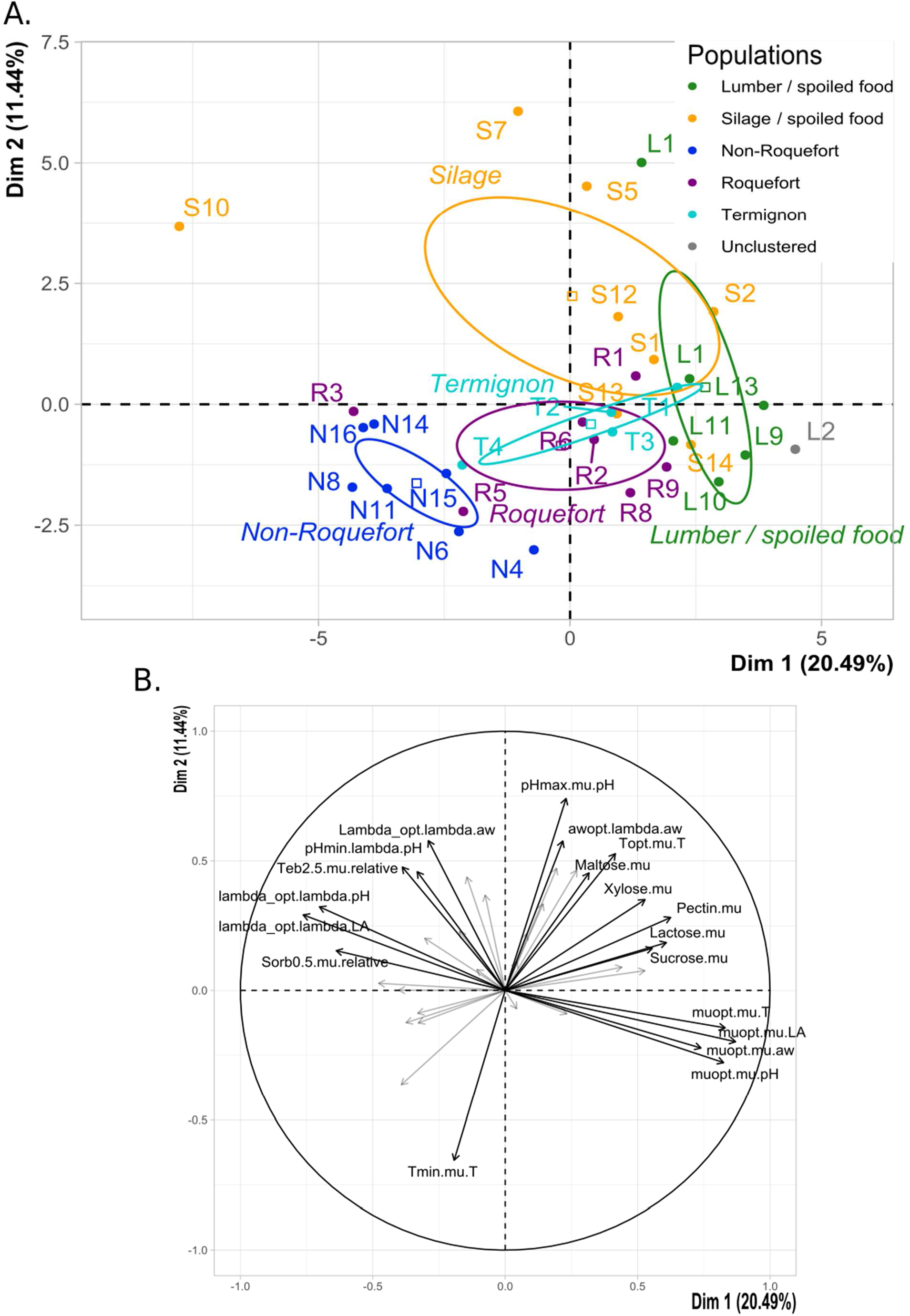
Principal component analysis (PCA) illustrating the phenotypic differences between *Penicillium roqueforti* populations based on growth response to temperature, water activity (salt), pH, various carbon sources (sucrose, glucose, lactose, galactose, maltose, cellobiose, xylose, starch, pectin and lactic acid) and to exposure to fungal inhibitors (lactic acid, potassium sorbate, tebuconazole and natamycin). **A.** Strains on the first two axes of the PCA. A confidence ellipse is drawn for each of the five populations. The percentage of variance explained by the axes are indicated. The same colour code is used as in the other figures: green for the lumber/spoiled food population, orange for the silage population, dark blue for the non-Roquefort cheese population, purple for the Roquefort cheese population and light blue for the Termignon cheese population. The strain IDs are provided in Suppl. Table 1. **B**. Association between the two PCA axes and the variables. Pectin.mu, lactose.mu, Maltose.mu, Xylose.mu, Sucrose.mu correspond to growth rate with pectin, lactose, maltose, xylose or sucrose as sole carbon sources, respectively. awopt.lambda.aw corresponds to optimal water activity (salt concentration minimising latency). Tmin.mu.T and Topt.mu.T corresponds to the minimal and optimal growth temperature; muopt.mu.T, muopt.mu.aw, muopt.mu.ph and muopt.mu.LA correspond the optimal parameter values for growth rate in terms of temperature, water activity (salt concentration), pH and lactic acid concentration, respectively. lambda.opt.lambda.aw, lambda.opt.lambda.pH and lambda.opt.lambda.LA, correspond to the optimal parameter values for inverse latency in terms of water activity (salt concentration), pH and lactic acid concentration, respectively. See Suppl. Fig. S1 for an illustration of parameter determination.

We included a negative control in each microplate (*i.e*. a well with only medium) for which the measures obtained were fitted with a polynomial model adapted to this type of data, *i.e*., with limited variation of signal. For each well, each fitted value for the negative control was subtracted to the value of the same time point obtained in inoculated wells in order to standardise the kinetics modelling across plates and conditions.

Using the output parameters of the primary modelling, we performed a secondary modelling for each of the studied parameters (*i.e*. temperature, pH, lactic acid concentration and water activity), to determine the cardinal values, *i.e*. the minimum, optimal and maximum values of these factors for growth, as well as the growth rate at the optimal value (*i.e*. the maximal growth rate across conditions, hereafter called “optimal growth rate” as typically done in growth rate modelling studies; see Figure 2A). We used cardinal models to fit the maximal growth rate and inverse latency (the inverse of the latency as this can be modelled with the same model as for the growth rate, in contrast to latency), to assess the impact of the tested abiotic factors on microbial growth using equations (2) to (5).

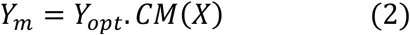

where CM(*X*) is the value returned by the cardinal model for *X*, the model being as in equation (3), (4) or (5) depending on the studied parameter *X*, *Y_opt_* corresponds to the optimal growth rate (RNU/hour) or optimal reciprocal latency (inverse of latency, λ^-1^_opt_) to model the effect of the various factors on the optimal growth rate or optimal inverse latency, noted *Y* in the equation.

To determine temperature and water activity cardinal values (*i.e*. minimum, optimal and maximum values for growth; Fig. 2A), we used the Rosso model (Rosso *et al*., 1993, 1995), largely used to model the effect of temperature and water activity on fungal growth (Dagnas & Membré, 2013), and presented in equation (3).

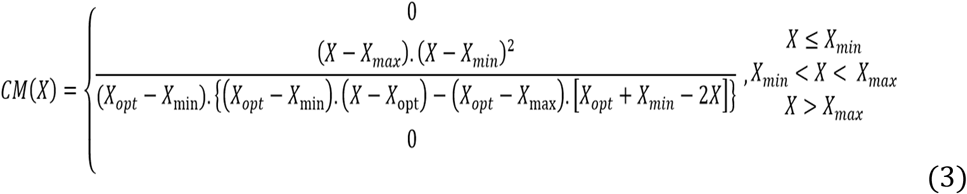

where *X* represents the studied parameter (temperature or water activity), and *X_min_, X_max_* and *X_opt_* are the minimum, maximum and optimal parameter values for growth, respectively (Fig. 2A). As the maximal water activity value is 1 (pure water), *a_w max_* was fixed at 1 (Pinon *et al*., 2004). As the growth rate was always lower than that measured in the maximal tested water activity value, *a_w opt_* was fixed at the highest *a_w_* value used, *i.e*. 0.996 (Savary et al. 2022). See Suppl. Fig. S1 for an illustration of the parameters.

For determining pH growth limits, we used the Presser model (Presser *et al*., 1998) presented in equation (4), that is the most appropriate when growth rate and latency are stable in a large range of pH as is the case in fungi:

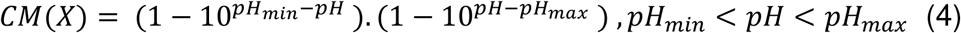

where pH_*min*_ and pH_*max*_ are the minimum and maximum pH values for which fungal growth is possible, respectively.

To model growth parameters as a function of lactic acid concentration, we applied the model developed by Le Marc et al. (2002), designed for estimating the minimum inhibitory concentration:

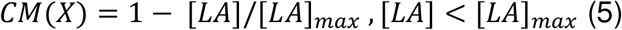

where [LA] represents the total concentration in lactic acid and [LA]*max* the maximum lactic acid concentration for fungal growth.

Growth rates and latencies obtained from primary modelling were fitted with these equations by minimising the sum of squared error (*nlinfit* function, MATLAB R2018b). Five percent confidence intervals were computed using the Jacobian matrix (*nlparci* function, MATLAB R2018b). We assessed how well our models predicted the data by computing the root mean square error (RMSE), measuring the error of a model in predicting quantitative data, and the determination coefficient (r^2^), measuring the proportion of variance in the growth parameter explained by the factor variation in the model. We kept only the values of the growth dynamics that could be modelled with sufficient confidence, hence strain replica values with coefficient r^2^ below 0.95 were excluded from further analyses. For population comparisons, we used a single value per strain and parameter, corresponding either to the mean of the replicates per strain (growth curve modelling) or to the secondary modelling outcome.

### Statistical analyses on phenotypes

In order to evaluate the impact of fungicides (lactic acid, potassium sorbate, tebuconazole and natamycin), controlling for intrinsic growth variations among strains, we computed relative latency and growth rate by dividing the measured growth values by a reference value for each strain. The reference values were the individual growth values obtained in PDB without fungicide.

Statistical analyses for testing differences in phenotypes between populations were performed using the R software (version 4.2.1, https://www.r-project.org/). For population comparisons, Shapiro-Wilk and Bartlett tests (package *rstatix*, R) were performed to assess normality and homoscedasticity of residuals in each population. If the data or the log-transformed data did not deviate from normality and homoscedasticity, populations were compared using ANOVA type I and Tukey tests were used as post-hoc tests. If the log-transformed and square-root transformed data significantly deviated from normality, a Kruskall-Wallis test was performed on raw data to compare populations, followed by Dunn tests as post-hoc tests.

To illustrate phenotypic population differentiation, we performed a principal component analysis (PCA) based on the phenotypes, *i.e*. the parameter values obtained in growth curves in all experiments using the R software (https://www.r-project.org/).

## Results

### Identification of a specific Termignon cheese population

After filtering, 190,387 SNPs were kept. The neighbour-net network (SplitsTree, Fig. 1A) revealed a novel *P. roqueforti* population, specific to Termignon cheeses, and most closely related to the non-Roquefort population (Fig. 1, Suppl. Table S2). The Termignon population displayed reticulations up to the tips of the network, suggesting that recombination may occur. The pairwise homology index tests in fact indicated signs of recombination in the Termignon population, as in the silage/spoiled food and lumber/soiled food populations (P<0.0001), but not in the Roquefort or non-Roquefort populations (P=0.99). Three strains in the Termignon population carried the MAT1-1 mating type (ESE00424, ESE00428 and ESE00685) while the fourth one carried the MAT1-2 mating type (ESE00426). A fifth strain had been isolated from a Termignon cheese (ESE00425) but actually belonged to the Roquefort genetic cluster.

The PCA based on SNPs (Fig. 1B), and the F_ST_ and d_XY_ values (Suppl. Table S2) showed a high differentiation of the non-Roquefort and Roquefort populations from other populations and one to each other, in agreement with the previous inference of strong divergence following independent domestication of these two lineages. The Termignon population again appeared differentiated from the other populations on the PCA, and the F_ST_ and d_XY_ values (Suppl. Table S2). The Termignon population appeared genetically closer to the non-Roquefort population on the splitstree but closer to the non-cheese populations on the PCA. The strong differentiation of the cheese populations from other populations in the PCA is likely due to the existence of a high number SNPs that are fixed for a nucleotide within the non-Roquefort population, and to a lesser extent within the Roquefort population, and that display different nucleotides in other populations (Suppl. Table S2), likely arising from bottlenecks and strong selection in these two cheese populations. The high percentage of identical and fixed alleles between the Termignon and the non-Roquefort populations (Suppl. Table S2) explains, on the other hand, that they are branched together in the neighbour-net network, *i.e*. showing that they are most closely related to each other. The silage/spoiled food and lumber/spoiled food had the highest diversity levels and shared polymorphism level (Suppl. Table S2), in agreement with the previous inference that they represent the most recent divergence in *P. roqueforti*.

The NGSAdmix analysis, which gives a probability of assignment to genetic clusters with a pre-defined number of clusters K, also supported the existence of five genetically differentiated clusters, with five well defined clusters at K=5, each including multiple strains assigned at 100% and with 70% of the runs finding this subdivision (Fig. 1C). At K=6, different subdivisions were inferred in the different runs, with balanced proportions for the various solutions, that inferred subclusters in either of the populations (Suppl. Fig. S1A and S1B). However, the splitstree and PCA (Fig. 1A and 1B) showed that the main level of genetic subdivision was at K=5. The second order rate of change in the likelihood (ΔK) peaked at K=4 (Suppl. Fig. S1C), indicating that the subdivision in four clusters is the strongest population subdivision level (the strongest increase in likelihood when increasing K by 1 was between K=4 and K=5). The neighbour-net network, the barplots, the PCA and the FST values (Fig. 1; Suppl. Table S1) nevertheless showed that the subdivision in five clusters is genuine and appeared biologically the most relevant regarding ecological niches. We therefore considered the subdivision into five clusters in the following.

A few strains displayed intermediate percentages of assignation at several clusters at K=5 and at higher K values. Such low percentages of assignation can result from admixture between groups or from low assignation power, for example because a few strains are differentiated from other strains within their cluster. The LCP06131 strain, for example, displayed intermediate assignment levels in barplots (Fig. 1C; Suppl. Fig. S1), but the splitstree showed that this strain is still much closer genetically to other strains in the Roquefort cluster than to other clusters, and without reticulations. Among the phenotyped strains, three strains showed intermediate percentages of assignment. The ESE00421 strain appeared intermediate between clusters in all analyses (splitstree, PCA and NGSAdmix; Fig. 1 and Suppl. Fig. S1), suggesting that it may be an admixed strain, *i.e*. resulting from hybridization between clusters. We therefore excluded this strain from the statistical tests on phenotype differences. Two other phenotyped strains, UBOCC-A-118017 and LCP06059, displayed intermediate assignment percentages, but fell in the middle of the silage/spoiled food cluster in the splitstree. The UBOCC-A-118017 strain appeared intermediate between the Termignon and silage/spoiled food clusters in the PCA but the LCP06059 strain clustered well in the silage group. Because of these assignment uncertainties, we tested differences between clusters with and without these two strains. We present below the statistical test results with the two strains included. Similar results were obtained when removing them from the dataset, unless specified otherwise.

The four Termignon strains carried the whole *CheesyTer* region in their genome, while only two (ESE00424 and ESE00685) harboured the *Wallaby* region. Only two strains (LCP04180 and LCP03969), both isolated from spoiled foods and belonging to the silage/spoiled food population, carried the *SORBUS* horizontally transferred region.

### *Penicillium roqueforti* populations exhibit distinct growth responses to abiotic factors

We tested whether growth behaviour differed between the five identified *P. roqueforti* populations in response to various abiotic factors. The principal component analysis (PCA) performed on the 33 phenotyped strains with the 39 inferred growth parameters (Suppl. Table S3) separated well the different *P. roqueforti* populations (Fig. 2). The first dimension separated the non-Roquefort and the lumber/spoiled food populations, the novel Termignon population being intermediate. This first PCA dimension was positively associated with the optimal growth rates obtained from secondary modelling in the temperature, water activity, pH and lactic acid concentration assays, and growth rate in lactose, pectin, xylose and sucrose, and negatively with minimum reciprocal latency and relative growth rate in potassium sorbate at 0.5 g.L^-1^. This suggests that the non-Roquefort population grows more slowly, is more tolerant to 0.5 g.L^-1^ potassium sorbate, has shorter minimal latency, and that the lumber / spoiled food population can better use lactose, pectin, xylose and sucrose for growth than the other populations.

The second PCA dimension separated the silage/spoiled food and cheese populations, the Termignon population once again appearing intermediate (Fig. 2). This second PCA dimension was positively associated with maximal pH, optimal water activity and optimal temperature, and negatively with minimal temperature. This suggests that cheese populations are less tolerant to higher pH and have lower optimal growth temperature than the silage/spoiled food population.

### The non-Roquefort population displays the highest salt and pH tolerance

Statistical tests confirmed a distinct growth behaviour of the non-Roquefort population compared to the other populations, revealing significant differences in terms of salt and pH tolerance. The populations indeed showed significant differences in the minimum water activity for growth (ANOVA Suppl. Table S4, becoming non-significant when the LCP06059 and UBOCC-A-118017 strains are excluded). *Post-hoc* tests did not detect any significant differences between pairs of populations (Fig. 3B, Suppl. Table S4). The means nevertheless indicated that the silage/spoiled food and non-Roquefort populations tended to tolerate lower minimum water activity (*i.e*. had higher salt tolerance) than lumber / spoiled food and Termignon populations (Fig. 3B, Suppl. Table S3). The non-Roquefort population was also characterised by a significantly lower optimal water activity for latency (mean of 0.993 for the shortest latency) than the Roquefort population (mean of 0.995 for the shortest latency, Suppl. Table S3). This means that the shortest latency for the non-Roquefort population was observed at a higher salt concentration (0.5% in NaCl) than for the Roquefort population (0.2% in NaCl). These findings support previous inference suggesting adaptation to highly salted environments in the non-Roquefort population.

**Figure 3:**
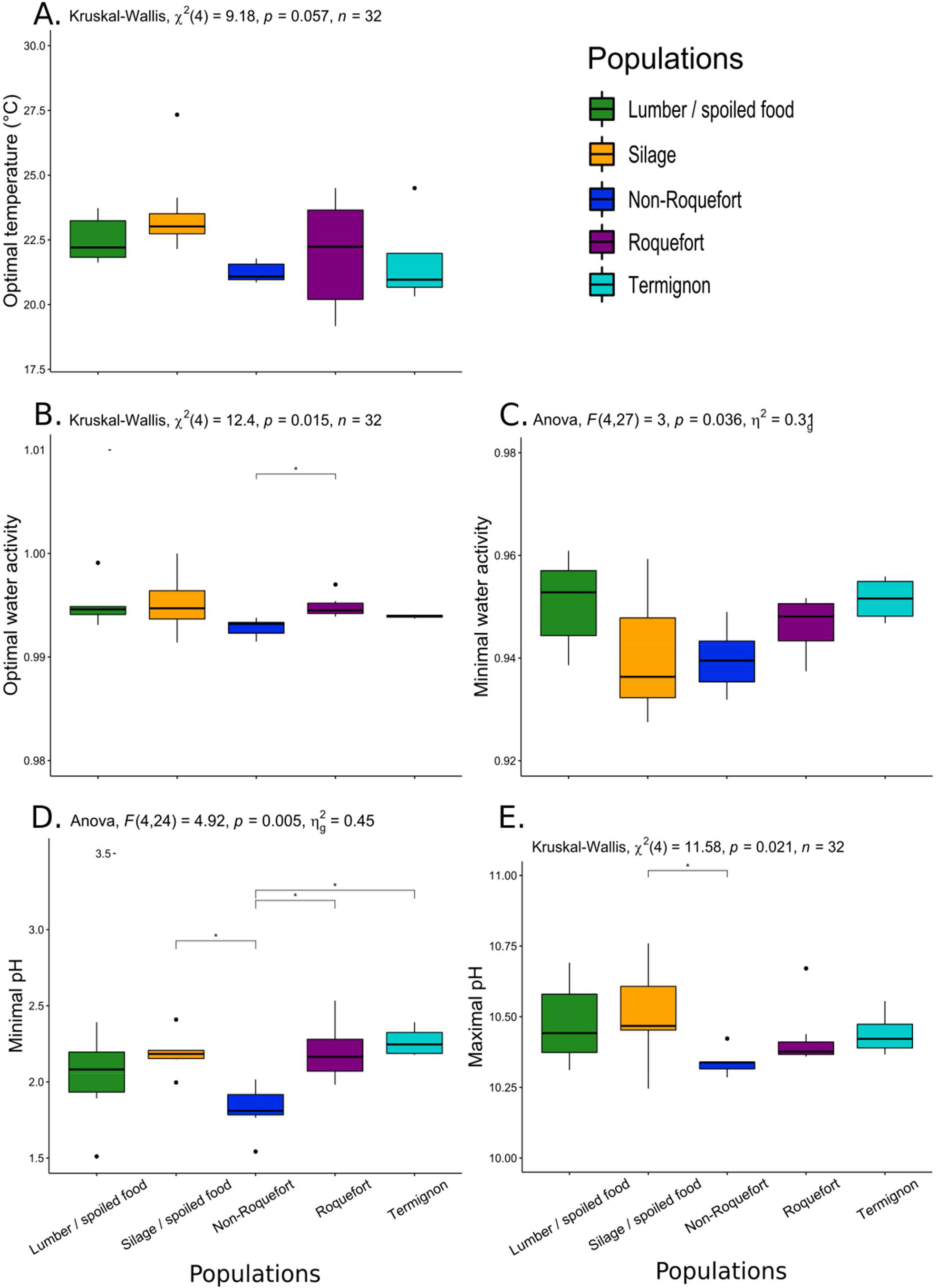
Growth parameters with differences between the *Penicillium roqueforti* five populations. The same colour code is used as in the other figures: green for the lumber/spoiled food population, orange for the silage population, dark blue for the non-Roquefort cheese population, purple for the Roquefort cheese population and light blue for the Termignon cheese population. The results of the global test for a population effect is given at the top of each panel. Pairwise significant differences are indicated by asterisks. The boxplots represent the median (centre line), the first quartile and third quartile (box bounds), the maximum and minimum excluding outlier points (whiskers), points being the outliers, *i.e*. with values either below the first quartile minus 1.5 fold the interquartile range or above the third quartile plus 1.5 fold the interquartile range. **A**. Optimal temperature, **B**. Optimal water activity, **C**. Minimal water activity (*i.e*. the fraction of available water for growth, which decreases for increasing salt concentrations), **D**. Minimal pH and **E**. Maximal pH.

The non-Roquefort population also tolerated more acidic pH, its mean minimal pH for growth being significantly lower than those for the silage/spoiled food, Roquefort and Termignon populations (mean pH_min_: 1.82 vs. 2.19, 2.20 and 2.27, respectively, Suppl. Table S3). The maximal pH for growth for the non-Roquefort population was also significantly lower than for the silage/spoiled food population (mean pH_max_: 10.34 vs. 10.51; Fig. 3D, Suppl. Table S3).

We detected a significant difference between populations in the minimal temperature for growth, post-hoc tests indicating a significantly lower minimal temperature for the Termignon population than for each of the two non-cheese populations (Fig. 2, Suppl. Table S4). However, the estimated minimal temperatures (T_min_) were all below 0°C, which has little biological or practical relevance in terms of cheese making, food storage or silage storage. Differences between populations were not significant for the other estimated cardinal temperature values, while there was a tendency for a lower temperature optimum for the non-Roquefort population compared to the silage population (Suppl. Table S4).

### Better capacity to use various carbohydrates for the silage/spoiled food and lumber/spoiled food populations

We also compared the ability of the different *P. roqueforti* populations to use various carbohydrates and lactic acid as carbon sources because cheese presents low concentrations in carbohydrates but a high concentration in lactic acid, in contrast to silage. The lumber/spoiled food population showed a significantly higher mean growth rate in potato dextrose broth than the non-Roquefort population (Fig. 4A and 4B Suppl. Table S4). We detected significant differences in growth rate between populations in lactose, sucrose, xylose and pectin (Fig. 4B, Suppl. Table S5). The lumber/spoiled food population indeed showed significantly faster growth than the non-Roquefort population in sucrose, xylose, and pectin (plant-derived sugars; Fig. 4B, Suppl. Table S5) and than the Roquefort population in lactose (dairy-derived sugar, Fig. 4B, Suppl. Table S5). The silage/spoiled food population grew significantly faster than the non-Roquefort population in pectin and than the non-Roquefort and Roquefort populations in lactose (Fig. 4B, Suppl. Table S5). The differences between the silage and non-Roquefort populations in terms of growth rate in pectin and lactose were not significant any more when removing the LCP06059 and UBOCC-A-118017 strains from the analysis. We did not detect any significant growth rate differences between populations for the other tested carbon sources (galactose, lactic acid, maltose or cellobiose, Fig. 4B, Suppl. Table S5).

**Figure 4:**
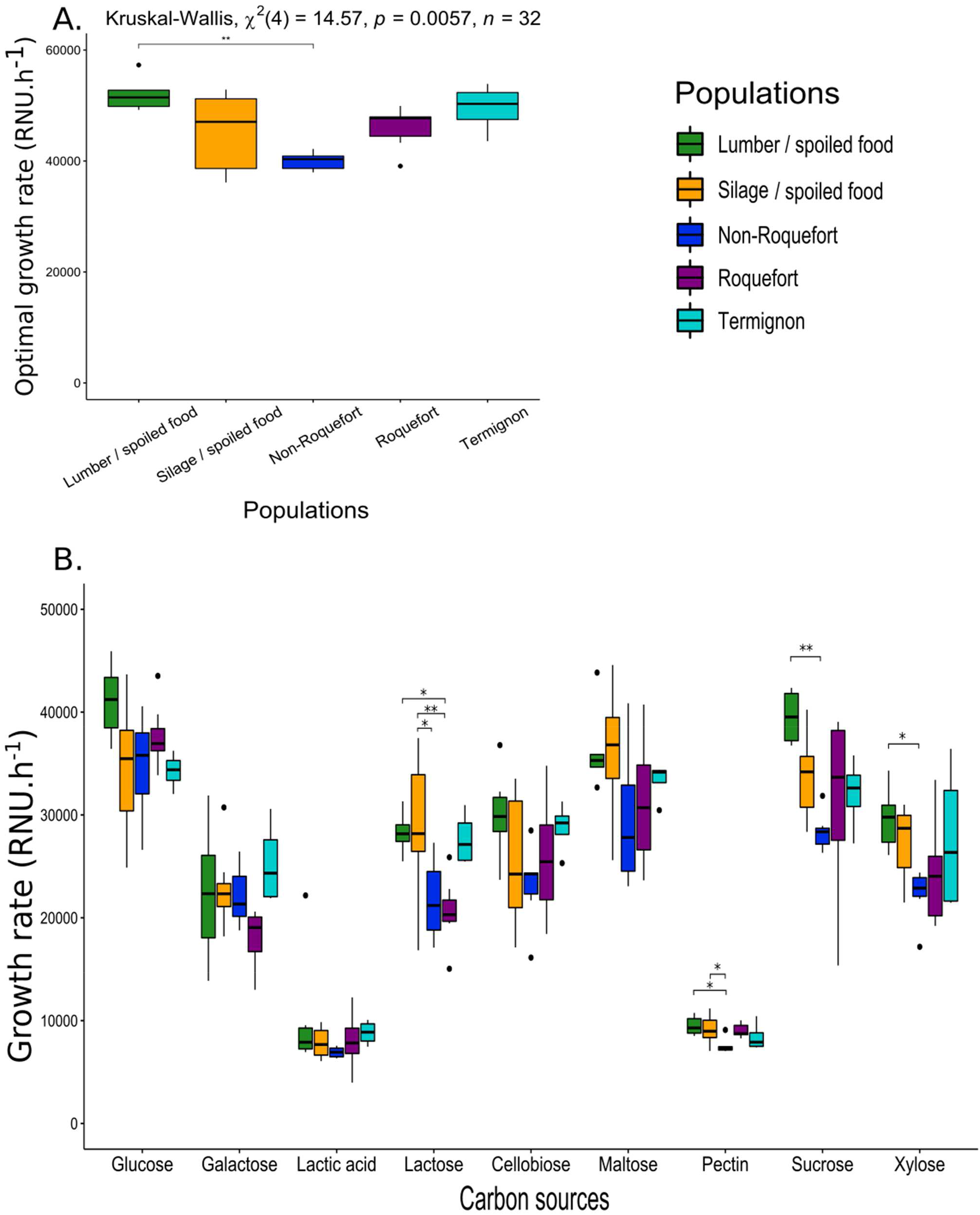
Differences in growth rate with various carbon sources between the *Penicillium roqueforti* populations. Pairwise significant differences are indicated by asterisks. The same color code is used as in the other figures: green for the lumber/spoiled food population, orange for the silage population, dark blue for the non-Roquefort cheese population, purple for the Roquefort cheese population and light blue for the Termignon cheese population. The boxplots represent the median (centre line), the first quartile and third quartile (box bounds), the maximum and minimum excluding outlier points (whiskers), points being the outliers, *i.e*. with values either below the first quartile minus 1.5 fold the interquartile range or above the third quartile plus 1.5 fold the interquartile range. **A**. Optimal growth rate, *i.e*. estimated growth rate at the optimal temperature, in potato dextrose broth, obtained from temperature secondary modelling. The results of the global test for a population effect is given at the top. **B**. Growth rate in minimal medium with glucose, galactose, lactic acid, lactose, cellobiose, maltose, pectin, sucrose and xylose as sole carbon sources.

### *Penicillium roqueforti* populations exhibit different growth responses to lactic acid exposure but no difference for other fungicides

As *P. roqueforti* populations can be exposed to various fungicides or fungal inhibitors depending on their ecological niches, we tested whether some of the *P. roqueforti* populations had higher tolerance than others. We detected significantly different behaviours between populations in response to the presence of lactic acid, a fungal inhibitor produced during lactic fermentation, with a higher tolerance of the non-Roquefort population (Fig. 5A). The non-Roquefort population growth rate was less impacted by lactic acid than other populations. Indeed, the maximal lactic acid concentration for growth was higher for the non-Roquefort population (mean of 1.05 M) than the other populations (means between 0.65 and 0.76 M, Fig. 5A, Suppl. Table S4); *Post-hoc* tests indicated that the difference between non-Roquefort and Roquefort population was significant. The non-Roquefort population showed higher relative growth rate at lactic acid concentrations ranging from 0.1 to 0.4 M (Fig. 5B, Suppl. Table S6) than other populations. This may result from an adaptation of the non-Roquefort population to the cheese environment where lactic acid is an important inhibitor. We detected no significant latency or growth rate differences between populations when exposed to the other fungicides, *i.e*. potassium sorbate and natamycin, used in the food industry, or tebuconazole, the main fungicide used on crops (Suppl. Table S6). Among the phenotypically tested strains, LCP04180, the only strain carrying the *SORBUS* region associated with sorbate-resistance, had a latency that was the least impacted by a sorbate concentration at 1 g.L^-1^ (Suppl. Table S3).

**Figure 5:**
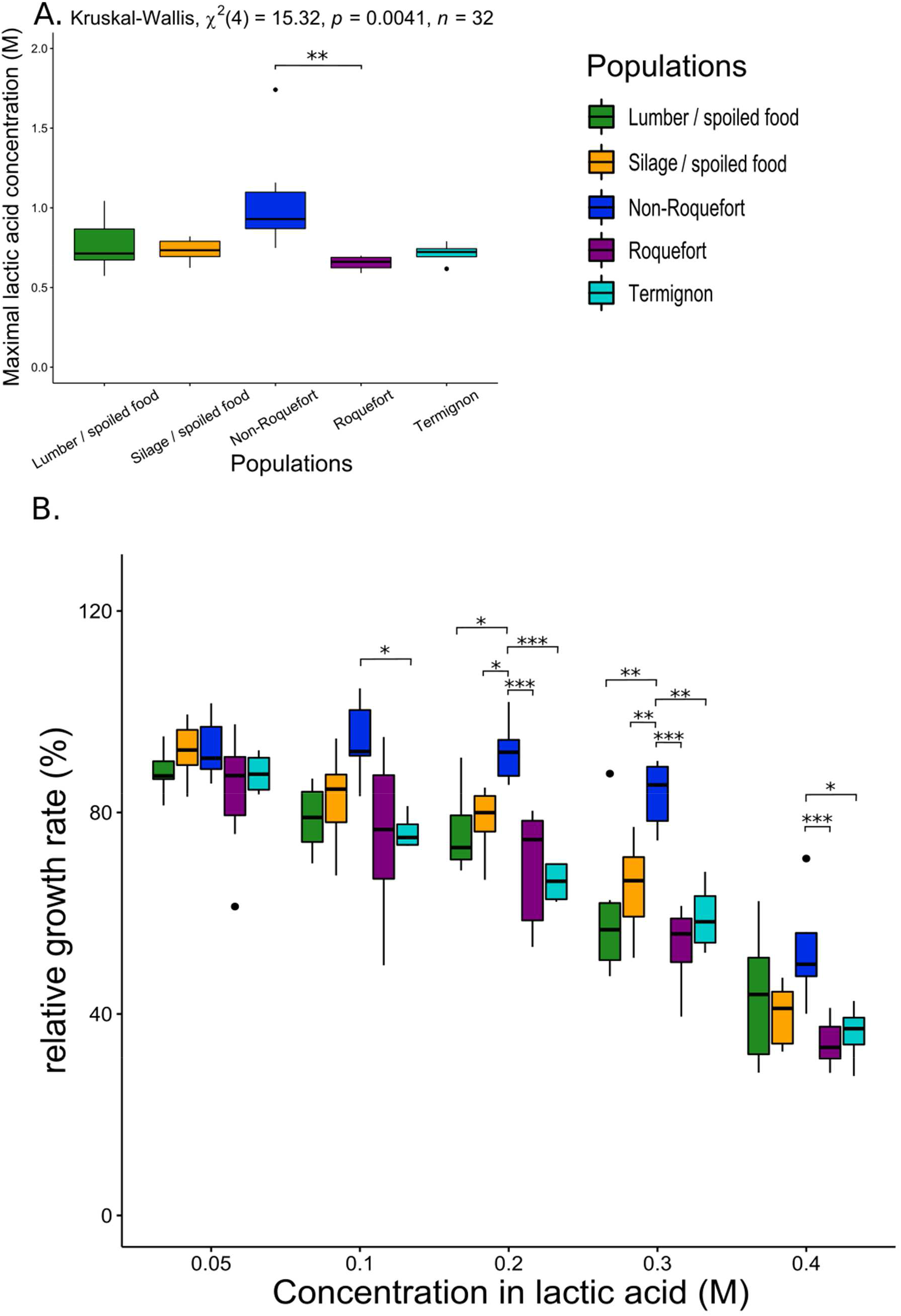
Impact of lactic acid on growth parameters between the *Penicillium roqueforti* populations. Pairwise significant differences are indicated by asterisks. The same color code is used as in the other figures: green for the lumber/spoiled food population, orange for the silage population, dark blue for the non-Roquefort cheese population, purple for the Roquefort cheese population and light blue for the Termignon cheese population.The boxplots represent the median (centre line), the first quartile and third quartile (box bounds), the maximum and minimum excluding outlier points (whiskers), points being the outliers, *i.e*. with values either below the first quartile minus 1.5 fold the interquartile range or above the third quartile plus 1.5 fold the interquartile range. **A**. Maximal lactic acid concentrations allowing growth for each of the five *Penicillium roqueforti* populations. The results of the global test for an effect of population maximal lactic acid concentration is given at the top. **B**. Relative growth rate with lactic acid concentration for each of the five *Penicillium roqueforti* populations. Relative latency and relative growth rate are growth rate and latency normalized by latency or growth rate without lactic acid.

## Discussion

In this study, we revealed a new *P. roqueforti* population, specific to Termignon cheeses, in which this blue mould is not inoculated. We detected differences between the cheese and non-cheese *P. roqueforti* populations in terms of growth response to various abiotic factors. The non-Roquefort cheese population appeared as the most phenotypically differentiated population, with higher tolerance to salt, more acidic pH and lactic acid. These results are in agreement with the previous suggestion that the non-Roquefort population has been subjected to strong selection for traits beneficial for cheese-making (Dumas *et al*., 2020). The Termignon population appeared phenotypically intermediate between cheese and non-cheese populations. The non-cheese populations show faster growth in various carbohydrates compared to the non-Roquefort and/or Roquefort populations, with faster growth of the lumber/spoiled food population in sucrose, xylose and pectin, and faster growth of the silage/spoiled food population in lactose and pectin. This suggests that the non-cheese populations are adapted to ecological niches that are rich in diverse carbohydrates, and particularly in plant-derived sugars.

### A specific *Penicillium roqueforti* population in Termignon cheeses

The Termignon strains, that spontaneously colonise cheeses in small-scale traditional cheese-making processes, formed a specific population, genetically differentiated from the four previously identified populations (Dumas *et al*., 2020). This population may be an unidentified, genuinely wild population, or a locally domesticated population. Its intermediate phenotypes between cheese and non-cheese populations suggest that it may have been selected for traits beneficial for cheese making, although less strongly than the other cheese populations. The fact that the Termignon strains are not cultivated or inoculated could let us think that they are not genuinely domesticated. They could however be feral strains deriving from an ancient cultivated *P. roqueforti* population, which would explain why the Termignon population is genetically most closely related to the non-Roquefort cheese population, carried the *CheesyTer* and for two strains the *Wallaby* regions (specific so far to the cheese non-roquefort population), and displayed phenotypes intermediate between cheese and non-cheese populations. They would thus be descendants of the domesticated population from which the non-Roquefort lineage would have been selected for and further improved, and thus remnants of a larger population from which most of the genetic diversity has been lost. We only isolated five strains from Termignon blue cheeses to date, including one that was assigned to the Roquefort population, so that we cannot rule out that strains belonging to other *P. roqueforti* populations may also colonise Termignon cheeses. However, the Termignon cheese production is made by only a handful of French farmers and at small scales (only a few hundred cheeses produced per year), so they may be mainly colonised by strains from the newly described population.

The identified Termignon population represents a potential new source of strain diversity for making blue cheeses, a highly valuable asset given that the Roquefort and the non-Roquefort populations have recently lost most of their diversity (Dumas *et al*., 2020). The discovery of this new population, with the two alternative mating types, paves the way for strain improvement, through the generation of offspring by crosses between populations, leveraging on the protocol developed to induce sexual reproduction in *P. roqueforti* (Ropars *et al*., 2016). This will generate novel genetic and phenotypic diversity for cheese making. All strains analysed so far from the non-Roquefort populations carried the MAT1-2 mating type, while most strains of the Roquefort populations harboured the MAT1-1 mating type. The presence of the two mating types in the Termignon population despite the low number of strains available reinforces the view that this population may be recombining, as suggested by the reticulation on the network and the Phi tests. This finding is of interest for future crosses with the two other cheese populations. New beneficial traits for cheese making could be obtained and sexual reproduction could counter-act the degeneration occurring in the clonal Roquefort and non-Roquefort lineages (Ropars *et al*., 2014, 2016).

### Specific traits in *Penicillium roqueforti* populations beneficial in their ecological niches

We detected signs of adaptation of the *P. roqueforti* populations to their ecological niches, in particular for the non-Roquefort cheese population and the non-cheese populations. A feature of cheese is its high salt content compared to other *P. roqueforti* ecological niches (wood, fruits and even bread). Salt has been traditionally used for taste enhancing and for its antimicrobial effect improving food conservation. Salt impairs growth of living organisms by decreasing water availability and through the toxicity of NaCl ions (Morin-Sardin *et al*., 2016; Venâncio *et al*., 2017). The domestication of fungi for cheese making may thus have led to salt tolerance as adaptation to the blue cheese environment, where salt concentration is 1.5 - 3.5% (El-Bakry, 2012; Hashem *et al*., 2014). In fact, we found here a higher salt tolerance in the non-Roquefort *P. roqueforti* population than in the other populations. This supports previous findings that showed higher salt tolerance for the non-Roquefort population based on growth experiments on malt agar and cheese-based media (Dumas *et al*., 2020). The Roquefort and Termignon cheese populations are less salt tolerant than the non-Roquefort population, which is consistent with previous observations showing that the non-Roquefort population exhibits more beneficial traits for commercial cheese production due to stronger selection (Dumas *et al*., 2020).

We found no particular tolerance for biologically relevant low temperature in the cheese populations, as in previous studies on other *Penicillium* cheese-making fungi (Ropars *et al*., 2020) or in other fungi used for cheese making, such as *Bisifusarium domesticum* and *Geotrichum candidum* (Savary *et al*., 2022; Bennetot *et al*., 2023). In the *S. cerevisiae* yeast, different modes of fermentation, at high versus low temperature, have yielded distinct lineages with different optimal temperatures (Baker *et al*., 2019).

The non-cheese populations also exhibited traits that may be beneficial in their ecological niche. The lumber/spoiled food population grew faster in potato dextrose broth and in various plant-derived carbohydrates (*e.g*. dextrose, sucrose, pectin and xylose). The silage/spoiled food population grew faster in pectin. These sugars are absent in cheese, in which proteins and lipids are instead dominant during ripening (Prieto *et al*., 2000). The faster growth of the non-cheese populations in plant-derived sugars may correspond to ancestral traits in *P. roqueforti*. Indeed, while no genuinely wild *P. roqueforti* population is known so far, it has been suggested that the ancestral population of the identified cheese and non-cheese populations could be a plant saprophyte or endophyte (Dumas *et al*., 2020). The carbohydrates used in our assays (*e.g*., xylose, dextrose, pectin and sucrose) represent important carbon sources in plants (Fukasawa & Matsukura, 2021). Important traits for wild populations can be lost in domesticated fungi due to the degeneration of unused traits or to drastic bottlenecks (Ropars & Giraud, 2022). Domestication in *P. roqueforti* could therefore have resulted in a fitness decrease in carbohydrate-rich media in the cheese populations. This is supported by the finding that the differences in growth rates of the lumber/spoiled food population were significant when compared to the most strongly domesticated cheese population, *i.e*. the non-Roquefort population. Cheese populations would have lost the ability to optimally use plant-derived sugars for growth and gained the ability to metabolise other nutrients (*e.g*. proteins and lipids). Cheese populations indeed have faster proteolytic activity and the non-Roquefort population has faster lipolysis than non-cheese populations in synthetic media (Dumas *et al*., 2020) and in the cheese environment (Caron *et al*., 2021). Alternatively, the ability to optimally use sugars may have been gained in the silage and the lumber/spoiled food populations as an adaptation to a new, anthropized ecological niche. The lack of identification of any undeniably wild *P. roqueforti* population precludes disentangling the two hypotheses.

The silage/spoiled food and lumber/spoiled food populations grew surprisingly faster in lactose than the Roquefort population. An explanation may be that, while galactose is abundant in milk as a constitutive hexose of lactose, it is also an important sugar in plant cell walls (Gangl & Tenhaken, 2016), which could lead to a selection in non-cheese populations for an efficient degradation of polymers containing galactose. Indeed, the same enzyme (β-galactosidase) is involved in the lysis of lactose and of plant cell wall hemicellulose polymers (*e.g*. xyloglucan and arabinogalactan proteins), which liberates galactose (Gangl and Tenhaken 2016; Fujita et al. 2019; Peña et al. 2004). Therefore, the faster growth of the non-cheese population in lactose may be a by-product of an adaptation to hemicellulose degradation. Alternatively, the non-cheese populations may have adapted to the use of lactose, as several strains isolated from spoiled food are also present in these populations (Dumas *et al*., 2020). In *S. cerevisiae*, the cheese population showed higher growth rate on galactose due to the acquisition, by horizontal gene transfer (HGT), of genes involved in the galactose import and degradation pathway (Legras *et al*., 2018). In our study, the non-Roquefort population did not exhibit higher efficiency for lactose use, despite the presence of genes involved in lactose import and degradation in the *Cheesyter* region acquired by HGT and only harboured by this population (Ropars *et al*., 2015).

Fungi can be exposed to various chemical biocides, in particular in crops and in food with preservatives. In laboratory experiments, recurrent exposure to biocides can lead to tolerance evolution (*i.e*. an increase in average minimum inhibitory concentrations), as shown for tebuconazole in *Aspergillus fumigatus* (Cui et al. 2019) and natamycin in *Aspergillus ochraceus, Fusarium oxysporum, Trichosporon asahii* and *Colletotrichum musae* (Streekstra et al. 2016). In *P. roqueforti*, sorbate resistance has been reported in some strains isolated from spoiled food, which has been attributed to the presence of a horizontally transferred region named *SORBUS* (Punt et al. 2022). Here, we studied the impact, on the growth of the different *P. roqueforti* populations, of chemical preservatives used in the food industry (natamycin and sorbate), a fungicide used for crop treatment (tebuconazole) and of a weak organic acid present in cheese (lactic acid resulting from lactose degradation by lactic acid bacteria). We did not detect any significant differences between the populations in terms of tolerance to potassium sorbate, tebuconazole and natamycin. This is consistent with our finding that a single silage strain carried the *SORBUS* region that provides sorbate resistance. This suggests that the selection for such tolerance is not strong in any of the ecological niches of *P. roqueforti* or that evolutionary constraints prevent adaptation.

Cheese and silage are rich in lactic acid due to the metabolism of lactic acid bacteria. We showed that the non-Roquefort population was more tolerant to lactic acid than the other populations, which may be an adaptive trait under cheese ripening conditions. In contrast, only a tendency for higher lactic acid tolerance was detected for the silage population, while silage can present higher concentrations in lactic acid than cheese (Marsili *et al*., 1981; Bevilacqua & Califano, 1989; Borreani & Tabacco, 2010). Organic acid tolerance has been reported in other domesticated fungi, for example tolerance to acetic acid, formic acid and levulinic acid in *S. cerevisiae* (Gallone *et al*., 2016). Tolerance to sulphite, a microbiocide classically used in wine-making, has also been reported in *S. cerevisiae* (Gallone *et al*., 2016; Legras *et al*., 2018) and in some wine-associated populations of the *Brettanomyces bruxellensis* yeast (Avramova *et al*., 2018).

## Conclusion

In conclusion, we report here new evidence for contrasting phenotypes between *P. roqueforti* populations, with beneficial traits for cheese making in the cheese populations (in particular higher tolerance to salt, acidic pH and lactic acid in the non-Roquefort population), which supports the inference of domestication in cheese fungi. In addition, we identified a novel *P. roqueforti* population, from non-inoculated French Termignon cheeses, with intermediate traits, substantial genetic diversity and the two mating types. This provides highly promising genetic resources for strain improvement, in addition to being a good model case to study domestication. We also found a more efficient use of various plant-derived carbon sources (in particular pectin, dextrose, xylose and/or lactose) in the silage/spoiled food and lumber/spoiled food populations, suggesting adaptation to anthropized niches, such as stored food and silage, and/or adaptation as saprophytes or endophytes. Among the three cheese populations, we found evidence for different stages of domestication, the non-Roquefort population showing the most differentiated traits, with phenotypes beneficial for cheese making, while the Termignon population appeared phenotypically intermediate between cheese and non-cheese populations. The non-Roquefort population also had the lowest genetic diversity, likely due to strong bottlenecks, which was not the case of the Roquefort population, certainly due to PDO requirements. Such gradual domestication syndromes have previously been reported in the soft-cheese making fungi from the *P. camemberti* clade (Ropars *et al*., 2020) and in the cheese-making fungus *Geotrichum candidum* (Bennetot *et al*., 2023). This suggests a protracted domestication process, with ancient, mild domestication, followed by more intensive selection in industrial times, as reported for example in maize (Allaby *et al*., 2008; Janzen & Hufford, 2016). The modern improvement process leads to a more drastic domestication syndrome, but also to worrying losses of diversity, threatening biodiversity conservation, jeopardising future strain improvement and leading to product uniformity (Vavilov, 2009). Our discovery of a new *P. roqueforti* population with higher diversity and contrasting phenotypes in Termignon cheeses is thus highly valuable for the sustainability of blue-cheese making and for strain improvement.

## Data accessibility

Genome accession numbers (Illumina reads and new assembly) to be added upon manuscript acceptance.

## Acknowledgments

This study was funded by the ANR-19-CE20-0002-02 Fungadapt and ANR-19-CE20-0006-01 Artifice ANR (French National Research Agency) grants. We thank Alodie Snirc and Stephanie Le Prieur for DNA extractions. We thank the GenoToul INRAe platform and the CNRS I2BC platform for genome sequencing.

## Author contributions

ECr performed experiments and analyses on phenotypes. JR, TG and AB collected the Termignon strains. AS performed DNA extraction and MinION sequencing. JR, TC and TG acquired the Termignon genomes. JR, JPV, AB and TC performed the genomic analyses. TG supervised the genomic analyses. JLJ, MC and ECo designed and supervised the experiments and analyses on phenotypes. AB contributed to statistical analyses. ECo, TG and JR obtained funding. ECr, TG and ECo wrote the first draft of the manuscript, with addition by JLJ, MC and JR. All authors revised the manuscript.

## Notes

### Competing Interest Statement

The authors have declared no competing interest.

