## Supplemental Table 1 for "A new cheese population in *Penicillium roqueforti* and adaptation of the five populations to their ecological niche"

**Table S1: *Penicillium roqueforti* strains used for population genetic analyses and phenotypic characterisation, with their IDs, assigned genetic cluster, sampling origin and date, genome accession number when available and the article in which they were published.** Lines in bold correspond to the strains used for phenotypic assays.

| Strain code | Short ID | Population | Sampling substrate | Sampling country | Sampling date | Genome accession number | Article source |
|---|---|---|---|---|---|---|---|
| **LCP06040** | **L1** | **Lumber / spoiled food** | **Drying Wood** | **France** | **1992** | **ERS1628921** | **Dumas et al, 2020** |
| LCP05419 | L3 | Lumber / spoiled food | Inner fridge wall | France | Unknown | ERS1628917 | Dumas et al, 2020 |
| LCP06039 | L4 | Lumber / spoiled food | Fruit Compote | URSS | 1973 | ERS1628920 | Dumas et al, 2020 |
| LCP04111 | L5 | Lumber / spoiled food | Wood | France | Unknown | ERS1628914 | Dumas et al, 2020 |
| LCP06060 | L6 | Lumber / spoiled food | Sulphite Liquor | Canada | Before 2014 | ERS1628924 | Dumas et al, 2020 |
| LCP06064 | L8 | Lumber / spoiled food | Mouldy baker's yeast | Denmark | Unknown | ERS1628925 | Dumas et al, 2020 |
| **LCP06037** | **L9** | **Lumber / spoiled food** | **Drying Wood** | **France** | **Unknown** | **ERS1628919** | **Dumas et al, 2020** |
| **UBOCC-A-117107** | **L10** | **Lumber / spoiled food** | **Apple** | **Kazakhstan** | **1974** | **To be added upon acceptance** | **This article** |
| **UBOCC-A-117110** | **L11** | **Lumber / spoiled food** | **Soil** | **Russia** | **1982** | **To be added upon acceptance** | **This article** |
| **UBOCC-A-117111** | **L12** | **Lumber / spoiled food** | **Apple** | **Kazakhstan** | **2007** | **To be added upon acceptance** | **This article** |
| **UBOCC-A-117112** | **L13** | **Lumber / spoiled food** | **Wood** | **United Kingdom** | **2007** | **To be added upon acceptance** | **This article** |
| LCP06138 | N2 | Non-Roquefort | Blue Cheese; Cambozola black label | Germany | Unknown | ERS1628936 | Dumas et al, 2020 |
| LCP06173 | N3 | Non-Roquefort | Blue Cheese; Roquefort Vernieres | France | Unknown | ERS1628937 | Dumas et al, 2020 |

| | | | | | | | |
|---|---|---|---|---|---|---|---|
| ESE00422 | N4 | Non-Roquefort | Feta cheese | Australia | 2014 | To be added upon acceptance | This article |
| LCP06134 | N6 | Non-Roquefort | Blue Cheese; Blue Sunshine | USA | Unknown | ERS1628932 | Dumas et al, 2020 |
| LCP06132 | N7 | Non-Roquefort | Blue Cheese; Stilton | United-Kingdom | Unknown | ERS1628930 | Dumas et al, 2020 |
| LCP00146 | N8 | Non-Roquefort | Blue Cheese; Roquefort | France | Unknown | ERS1628909 | Dumas et al, 2020 |
| LCP06133 | N9 | Non-Roquefort | Blue Cheese; Roche Montagne | France | 2013 | ERS1628931 | Dumas et al, 2020 |
| LCP06135 | N10 | Non-Roquefort | Blue Cheese; Bath Blue Organic | England | Unknown | ERS1628933 | Dumas et al, 2020 |
| LCP06129 | N11 | Non-Roquefort | Blue Cheese; Crémeux du Puy | France | Unknown | ERS1628927 | Dumas et al, 2020 |
| UBOCC-A-117091 | N14 | Non-Roquefort | Subglacial water | Norway | 2006 | To be added upon acceptance | This article |
| UBOCC-A-117092 | N15 | Non-Roquefort | Blue Cheese; Bleu du Vercors Sassenage | France | 2016 | To be added upon acceptance | This article |
| UBOCC-A-118016 | N16 | Non-Roquefort | Irrigation water | France | 2016 | To be added upon acceptance | This article |
| ESE00424 | T1 | Termignon | Blue Cheese; Bleu de Termignon | France | 2018 | To be added upon acceptance | This article |
| ESE00426 | T2 | Termignon | Blue Cheese; Bleu de Termignon | France | 2018 | To be added upon acceptance | This article |
| ESE00428 | T3 | Termignon | Blue Cheese; Bleu de Termignon | France | 2018 | To be added upon acceptance | This article |
| ESE00685 | T4 | Termignon | Blue Cheese; Bleu de Termignon | France | 2015 | To be added upon acceptance | This article |
| ESE00170 | R1 | Roquefort | Peach | Unknown | 2018 | To be added upon acceptance | This article |
| LCP06136 | R2 | Roquefort | Blue Cheese; Great Hill Blue | USA | Unknown | ERS1628934 | Dumas et al, 2020 |
| ESE00425 | R3 | Roquefort | Blue Cheese; Bleu de Termignon | France | 2018 | To be added upon acceptance | This article |
| LCP00148 | R4 | Roquefort | Brewery environment | Unknown | Unknown | ERS1628910 | Dumas et al, 2020 |

| | | | | | | | |
|---|---|---|---|---|---|---|---|
| **LCP02939** | **R5** | **Roquefort** | **Brioche under plastic wrap** | **Unknown** | **Unknown** | **ERS1628911** | **Dumas et al, 2020** |
| **LCP06130** | **R6** | **Roquefort** | **Blue Cheese; Gorgonzola** | **Italy** | **2013** | **ERS1628928** | **Dumas et al, 2020** |
| LCP04157T | R7 | Roquefort | Blue Cheese; Roquefort | USA | 1987 | ERS1628915 | Dumas et al, 2020 |
| **UBOCC-A-113022** | **R8** | **Roquefort** | **Blue Cheese; Roquefort** | **France** | **2013** | **To be added upon acceptance** | **This article** |
| **UBOCC-A-116029** | **R9** | **Roquefort** | **Atmosphere cave** | **France** | **2016** | **To be added upon acceptance** | **This article** |
| LCP06128 | R12 | Roquefort | Blue Cheese; Roquefort | France | Unknown | ERS1628926 | Dumas et al, 2020 |
| LCP06131 | R13 | Roquefort | Blue Cheese; Roquefort | France | Unknown | ERS1628929 | Dumas et al, 2020 |
| LCP06137 | R14 | Roquefort | Blue Cheese; Roquefort Vernieres | France | Unknown | ERS1628935 | Dumas et al, 2020 |
| **LCP06059** | **S1** | **Silage / spoiled food** | **Silage** | **Netherlands** | **Unknown** | **ERS1628923** | **Dumas et al, 2020** |
| **LCP04180** | **S2** | **Silage / spoiled food** | **Strawberry sorbet** | **France** | **1997** | **ERS1628916** | **Dumas et al, 2020** |
| LCP03969 | S3 | Silage / spoiled food | Fruit Compote | France | 1996 | ERS1628913 | Dumas et al, 2020 |
| **LCP06635** | **S5** | **Silage / spoiled food** | **Silage** | **Belgium** | **2006** | **To be added upon acceptance** | **This article** |
| **LCP06667** | **S7** | **Silage / spoiled food** | **Silage** | **Belgium** | **2007** | **To be added upon acceptance** | **This article** |
| LCP06043 | S8 | Silage / spoiled food | Silage | Belgium | Unknown | ERS1628922 | Dumas et al, 2020 |
| **LCP06679** | **S10** | **Silage / spoiled food** | **Silage** | **Belgium** | **2007** | **To be added upon acceptance** | **This article** |
| LCP06036 | S11 | Silage / spoiled food | Cork Wine Bottle | Belgium | Unknown | ERS1628918 | Dumas et al, 2020 |
| **UBOCC-A-117102** | **S12** | **Silage / spoiled food** | **Silage** | **Belgium** | **2011** | **To be added upon acceptance** | **This article** |
| **UBOCC-A-117291** | **S13** | **Silage / spoiled food** | **Silage** | **France** | **2017** | **To be added upon acceptance** | **This article** |

| | | | | | | | |
|---|---|---|---|---|---|---|---|
| **UBOCC-A-118017** | **S14** | **Silage / spoiled food** | **Flour** | **France** | **2016** | **To be added upon acceptance** | **This article** |
| **ESE00421** | **L2** | **Unassigned** | **Bread** | **France** | **2019** | **To be added upon acceptance** | **This article** |

All strains are in compliance with the Nagoya protocol.
