## Supplemental Table 2 for "A new cheese population in *Penicillium roqueforti* and adaptation of the five populations to their ecological niche"

**Supplementary Table S2: Statistics of diversity and of differentiation between the five *Penicillium roqueforti* populations: non-Roquefort cheese (nonRoq), Roquefort cheese (Roq), Termignon cheese (Termignon), lumber/spoiled food (Lumber) and silage/spoiled food (Silage).** The strain ESE00421 was not included for these computations. **A**. Number of single nucleotide polymorphisms (SNPs) between pairs of populations, that are either fixed (fixed in each population and different between the two populations), identical (fixed and identical between the two populations), private (fixed in one population, polymorphic in the other population) and shared (polymorphic with shared alleles in the two populations). The highest number per column is in bold. **B.** Indices of diversity within populations (Pi and Whatterson's theta) and of differentiation between populations ($F_{ST}$ and $d_{XY}$).

**A**

| **Pair of populations** | **fixed** | **Identical** | **private** | **shared** | **Total** |
|---|---|---|---|---|---|
| nonRoq vs Lumber | 19006 | 918485 | 79755 | 3279 | 102040 |
| Roq vs Lumber | 6595 | 924267 | 68484 | 21179 | 96258 |
| Roq vs nonRoq | 28756 | 949448 | 40619 | 1702 | 71077 |
| Silage vs Lumber | 949 | 878760 | 87491 | 53325 | 141765 |
| Silage vs nonRoq | 8806 | 890465 | 115186 | 6068 | 130060 |
| Silage vs Roq | 3860 | 890296 | 100887 | 25482 | 130229 |
| Termignon vs Lumber | 10499 | 910304 | 81740 | 17982 | 110221 |
| Termignon vs nonRoq | 20645 | 952358 | 44159 | 3363 | 68167 |
| Termignon vs Roq | 19442 | 936511 | 53730 | 10842 | 84014 |
| Termignon vs Silage | 4447 | 885072 | 103299 | 27707 | 135453 |

**B**

| | pi | Watterson's theta |
|---|---|---|
| **non Roquefort** | 1,15E-04 | 1,21E-04 |
| **Roquefort** | 4,03E-04 | 4,07E-04 |
| **Termignon** | 8,69E-04 | 8,46E-04 |
| **Lumber-Spoiled food** | 1,06E-03 | 9,70E-04 |
| **Silage-Spoiled food** | 1,60E-03 | 1,58E-03 |

| **Fst** | **Termignon** | **Roquefort** | **Lumber-Spoiled food** | **Silage-Spoiled food** |
|---|---|---|---|---|
| **non Roquefort** | 0,69 | 0,85 | 0,71 | 0,57 |
| **Termignon** | | 0,64 | 0,51 | 0,36 |
| **Roquefort** | | | 0,52 | 0,4 |
| **Lumber-Spoiled food** | | | | 0,25 |

| **dxy** | **Termignon** | **Roquefort** | **Lumber-Spoiled food** | **Silage-Spoiled food** |
|---|---|---|---|---|
| **non Roquefort** | 0,00162 | 0,00171 | 0,00197 | 0,00202 |
| **Termignon** | | 0,00181 | 0,00195 | 0,00198 |
| **Roquefort** | | | 0,00151 | 0,00168 |
| **Lumber-Spoiled food** | | | | 0,00177 |
