## Supplemental Figure 1 for "A new cheese population in *Penicillium roqueforti* and adaptation of the five populations to their ecological niche"

**Supplementary Figure 1: Schematic representation of the experiments performed for testing the impact of different parameters on the growth of the five populations of the blue cheese fungus *Penicillium roqueforti*.** Growth dynamics was measured by laser nephelometry (cell density in liquid), then a primary modeling was applied to fit growth curves and estimate growth parameters (latency, growth rate and maximal growth); then, a secondary modeling was performed to estimate cardinal values (*i.e.* minimal, maximal and optimal values of the considered parameter for growth and the optimal growth, *i.e.*, the growth rate at the optimal value, corresponding to the maximal growth rate.

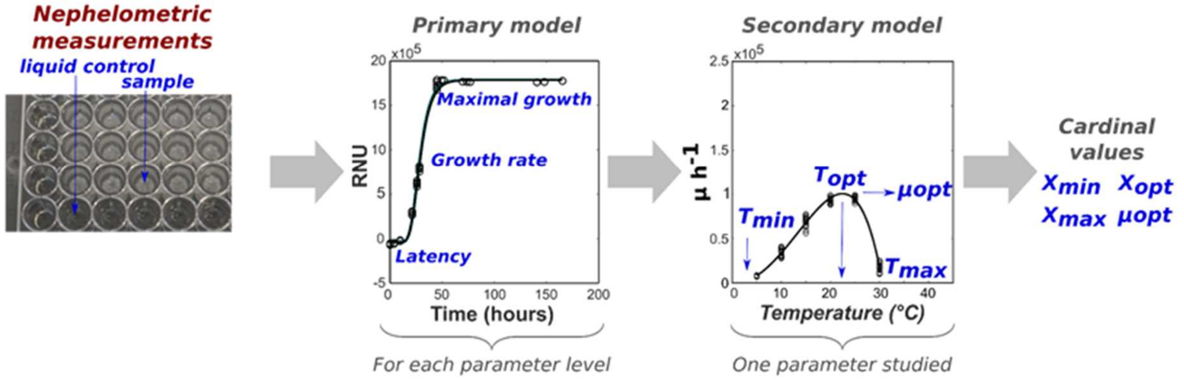
