## Supplemental Figure 2 for "A new cheese population in *Penicillium roqueforti* and adaptation of the five populations to their ecological niche"

**Supplementary Figure 2: Population subdivision inferred with NGSadmix. A.** Population subdivision inferred for K=2 to 6. Colored bars represent the coefficients of membership in the K gene pools based on genomic data. Each bar represents a strain, its name being indicated at the bottom of the figure. The ID of the strains used for phenotyping are in bold. The barplots represent, for each K value, the solution inferred in the highest percentage of the 100 runs and this percentage is indicated at the top. **B.** Different solutions inferred at K=6, with their percentage among the 100 runs given at the top. **C.** Second order rate of change in the likelihood ($\Delta K$, at left) and log of the likelihood across K values.

**A**

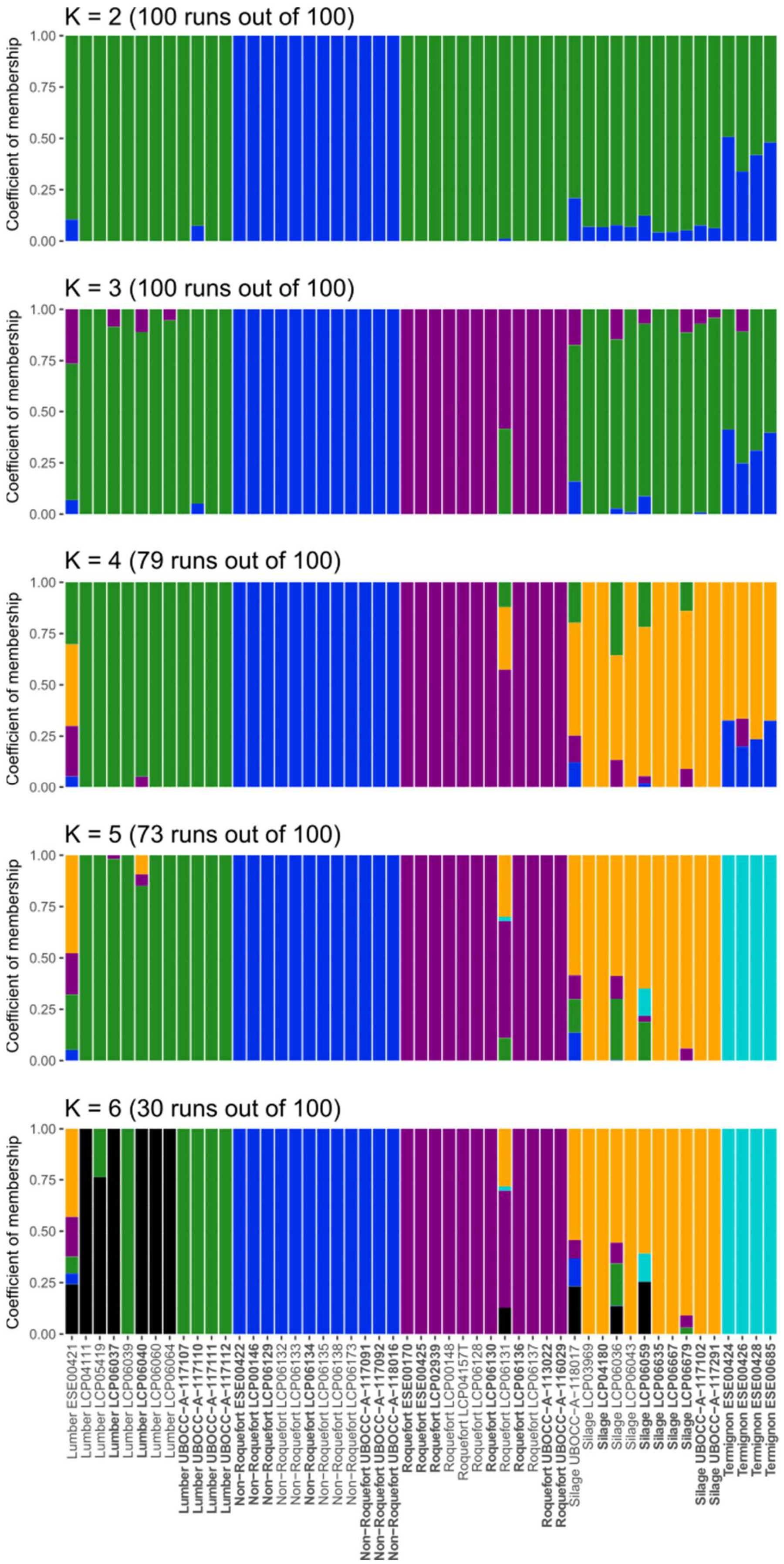
K = 2 (100 runs out of 100)
K = 3 (100 runs out of 100)
K = 4 (79 runs out of 100)
K = 5 (73 runs out of 100)
K = 6 (30 runs out of 100)
Coefficient of membership
0.00
0.25
0.50
0.75
1.00
Lumber ESE00421
Lumber LCP04111
Lumber LCP05419
Lumber LCP06037
Lumber LCP06039
Lumber LCP06040
Lumber LCP06060
Lumber LCP06064
Lumber UBOCC-A-117107
Lumber UBOCC-A-117110
Lumber UBOCC-A-117111
Lumber UBOCC-A-117112
Non-Roquefort ESE00422
Non-Roquefort LCP00146
Non-Roquefort LCP06129
Non-Roquefort LCP06132
Non-Roquefort LCP06133
Non-Roquefort LCP06134
Non-Roquefort LCP06135
Non-Roquefort LCP06138
Non-Roquefort LCP06173
Non-Roquefort UBOCC-A-117091
Non-Roquefort UBOCC-A-117092
Non-Roquefort UBOCC-A-118016
Roquefort ESE00170
Roquefort ESE00425
Roquefort LCP02939
Roquefort LCP00148
Roquefort LCP04157T
Roquefort LCP06128
Roquefort LCP06130
Roquefort LCP06131
Roquefort LCP06136
Roquefort LCP06137
Roquefort UBOCC-A-113022
Roquefort UBOCC-A-116029
Silage UBOCC-A-118017
Silage LCP03969
Silage LCP04180
Silage LCP06036
Silage LCP06043
Silage LCP06059
Silage LCP06635
Silage LCP06667
Silage LCP06679
Silage UBOCC-A-117102
Silage UBOCC-A-117291
Termignon ESE00424
Termignon ESE00426
Termignon ESE00428
Termignon ESE00685

**B**

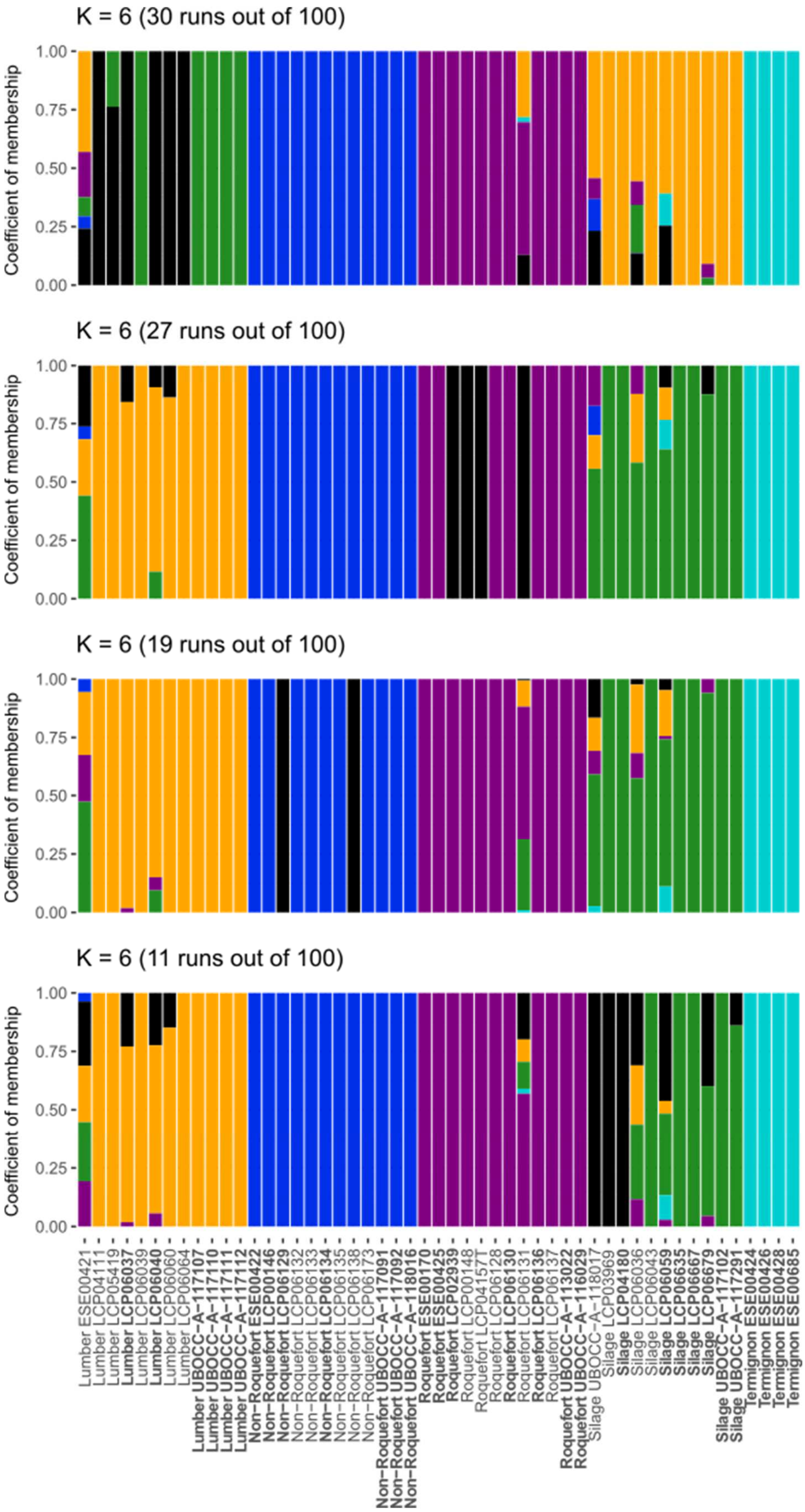
K = 6 (30 runs out of 100)
K = 6 (27 runs out of 100)
K = 6 (19 runs out of 100)
K = 6 (11 runs out of 100)
Coefficient of membership
1.00
0.75
0.50
0.25
0.00
Lumber ESE00421
Lumber LCP04111
Lumber LCP05419
Lumber LCP06037
Lumber LCP06039
Lumber LCP06040
Lumber LCP06060
Lumber LCP06064
Lumber UBOCC−A−117107
Lumber UBOCC−A−117110
Lumber UBOCC−A−117111
Lumber UBOCC−A−117112
Non−Roquefort ESE00422
Non−Roquefort LCP00146
Non−Roquefort LCP06129
Non−Roquefort LCP06132
Non−Roquefort LCP06133
Non−Roquefort LCP06134
Non−Roquefort LCP06135
Non−Roquefort LCP06138
Non−Roquefort LCP06173
Non−Roquefort UBOCC−A−117091
Non−Roquefort UBOCC−A−117092
Non−Roquefort UBOCC−A−118016
Roquefort ESE00170
Roquefort ESE00425
Roquefort LCP02939
Roquefort LCP00148
Roquefort LCP04157T
Roquefort LCP06128
Roquefort LCP06130
Roquefort LCP06131
Roquefort LCP06136
Roquefort LCP06137
Roquefort UBOCC−A−113022
Roquefort UBOCC−A−116029
Silage UBOCC−A−118017
Silage LCP03969
Silage LCP04180
Silage LCP06036
Silage LCP06043
Silage LCP06059
Silage LCP06635
Silage LCP06667
Silage LCP06679
Silage UBOCC−A−117102
Silage UBOCC−A−117291
Termignon ESE00424
Termignon ESE00426
Termignon ESE00428
Termignon ESE00685

**C**

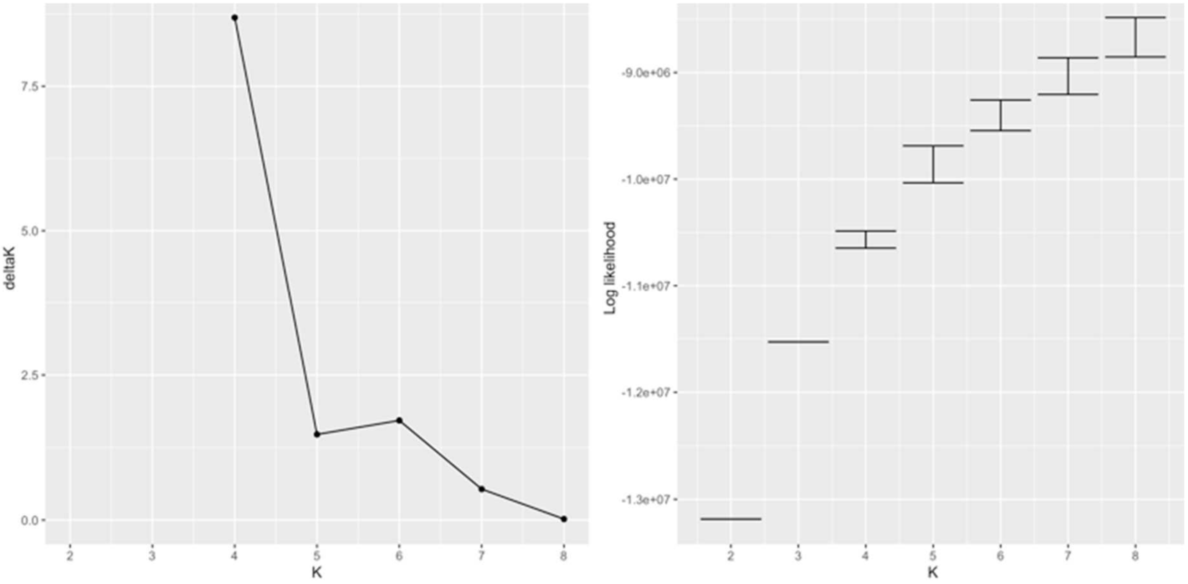
deltaK
0.0
2.5
5.0
7.5
2
3
4
5
6
7
8
K
Log likelihood
-9.0e+06
-1.0e+07
-1.1e+07
-1.2e+07
-1.3e+07
2
3
4
5
6
7
8
K
